# Aging changes the underlying mechanism of JAK2 modulation in neutrophil function

**DOI:** 10.1101/2025.04.21.649782

**Authors:** Jacob W. Feldmann, Matthew Kays, Farrah McGinnis, Emily Herron, Nurullah Sati, Clara Woods, Aminata P. Coulibaly

## Abstract

Janus Kinase 2 (JAK2) has been linked to various neutrophil functions, but the intracellular mechanisms underlying its modulation are unknown. Neutrophils are essential cells for host defense. Neutrophil effector functions include migration, reactive oxygen species (ROS) production, degranulation, and neutrophil extracellular trap (NET) formation. The goal of this study was to elucidate the signaling mechanism through which JAK2 modulates neutrophil function and the effect of aging on this pathway. We hypothesized that JAK2-mediated modulation changes the molecular mechanisms associated with neutrophil function in an age- and sex-dependent manner. Neutrophils from young (3 months) and aged (22+ months), male and female C57BL/6J mice were isolated, treated with a JAK2 inhibitor (AZD1480) or a pan-JAK inhibitor (Baricitinib), and stimulated with Phorbol 12-myristate 13-acetate (PMA). Functional assays were conducted to assess migration, ROS production, degranulation, NETosis, and metabolism. Mass spectrometry and Luminex assays provided proteomic and cytokine profiles. Our data show that JAK2 promotes migration via membrane composition and actin remodeling, with age-dependent shifts in chemokine secretion. JAK2 indirectly affects NETosis by modulating IL-1 signaling and ROS production. It also primes ROS production by altering NADPH oxidase components. In young neutrophils, JAK2 influences degranulation through actin remodeling, while aged neutrophils display impaired granule release. Metabolically, JAK2 enhances pentose phosphate pathway activity in young neutrophils and decreases glycogen breakdown in aged cells. These findings reveal mechanisms by which JAK2 modulates neutrophil function and suggest age-specific therapeutic targeting in inflammatory diseases.

## Introduction

Neutrophils are the most abundant innate immune cell in the mammalian body. They are the first immune cells to be recruited to sites of inflammation, injury, and infection [1-3]. Neutrophils comprise 40-80% of all white blood cells in the mammalian system [4]. Neutrophils have short half-lives, leading to a high turnover rate. Due to their importance in the inflammatory response and abundance in circulation, it is important to understand the biology of this immune cell type. Neutrophil effector functions are critical to their activity. These effector functions include production of cytokines to mediate pro- or anti-inflammatory environments, removal of debris and damaged cells through phagocytosis. Additionally, neutrophils produce reactive oxygen species (ROS), in order to aid in cellular breakdown or cell signaling [5], and neutrophil extracellular traps (NETs) [6]. All these functions are modulated by various intracellular signaling cascades initiated by various ligands and receptor interactions at the cell plasma membrane. It is becoming evident that various physiological factors affect neutrophil function, including sex and aging. Three recent studies have shown sex differences in neutrophil activation and omics-landscape (transcription, proteomic, metabolomic and lipidomic) in response to stimuli [7-9]. In circulating human neutrophils, female neutrophils have heightened type I interferon (IFN) and Toll-like receptor responses, and distinct maturation patterns, while male neutrophils have increased mitochondrial metabolism in the presence of estradiol (in a study of males with Klinefelter syndrome [XXY] [7]). Circulating human male and female neutrophils differ in mRNA translation efficiency, protein abundance, and phosphorylation levels [8]. Single-cell RNA sequencing of bone marrow-derived mouse neutrophils reveals distinct sex-specific transcriptional profiles across neutrophil subpopulations. Female neutrophils exhibit transcriptional signatures associated with functional responses, including motility, IFN-γ signaling, and VEGF signaling, whereas male neutrophils display a profile enriched for proliferative pathways, such as cell cycle checkpoints, M-phase progression, and mitotic pro-metaphase regulation [9]. This study also shows that this transcriptional sexual dimorphism persists with age (4 months old vs. 20 months old) in neutrophils [9]. A study using human bone marrow from young (20-30 years of age) and aged (70-80 years of age) healthy volunteers showed blunted response to G-CSF with age [10]. In mice, circulating neutrophil numbers increase with age (with male neutrophils increasing at a faster rate compared to females), in a non-disease state [11]. These data suggest that age directly affects neutrophil response to stimuli, whether this is due to a change in intracellular integration of signals is unknown.

The JAK/STAT pathway is a ubiquitous signaling mechanism across various cell types, activated by extracellular stimuli like inflammatory cues (cytokines/chemokines) and growth factors [12-17]. There are four characterized JAK isoforms, JAK1, JAK2, JAK3, and TYK2. The activation of each JAK protein elicits different downstream effector proteins, which lead to different functional outcomes [18-21]. Our current understanding of JAK/STAT signaling in neutrophils is limited. JAK2 activation was shown to activate neutrophils, regulate migration [22], increase proliferation of neutrophil progenitor cells [23, 24], and enhance neutrophil fungal killing through the release of elastase, MMP9, and ROS [25]. In this project we aimed to elucidate whether age and sex affect JAK modulation of neutrophil function. We hypothesized that age and sex change the intracellular pathway used by JAK2 to modulate neutrophil function.

Our data show that aging abrogates the direct effect JAK2 activation has on neutrophil function. JAK2 activation is required for intracellular signaling cascades that underlie migration and degranulation. However, the release of chemokines is differentially regulated in young and aged neutrophils. Even with no direct effect on ROS production in both young and aged neutrophils, JAK2 regulates various protein components of the NADPH oxidase (NOX) complex. Similarly, JAK2 has no direct effect on NETosis in young and aged mice but has diverse age-related effects on critical NETosis proteins like neutrophil elastase (NE) and myeloperoxidase (MPO). Interestingly, JAK2 signaling plays a critical role in glucose shuffling in young neutrophils, while only influencing glycogen breakdown in aged neutrophils. No sex differences were observed in JAK2 modulation of neutrophil function. This study provides evidence of various cellular mechanisms in neutrophils that are modulated by JAK2 activation in an age dependent manner.

## Results

Neutrophils undergo an aging process after release from the bone marrow [26]. The length of temporal aging is species dependent. For example, mouse neutrophils have a lifespan of approximately 18 hours and those from humans 5.4 days [27]. Beside the temporal age of neutrophils [26], the biological age of the host influences the grade of cells released from the bone marrow. In this study, we investigate the effect of biological aging on neutrophil function under normal conditions. For the remainder of this article, the term ‘young neutrophils’ refers to neutrophils isolated from young mice (3 months) and ‘aged neutrophils’ to neutrophils isolated from aged mice (22+ months). To control for the temporal age all isolation of neutrophils were conducted at the same time each day. It is well established that Phorbol 12-myristate 13-acetate (PMA) modulates neutrophil function by activating various components of the Protein Kinase C (PKC) family of proteins. Through its effect on PKC-α, -βI, -βII, and -*δ* [28], PMA induces neutrophil migration [29], NOX formation, ROS production, and NETosis [30, 31]. To determine which neutrophil functions require JAK signaling, neutrophils from young and aged mice were exposed to a pan-JAK inhibitor (Baricitinib [32]) or a JAK2 selective inhibitor (AZD1480 [33]) followed by PMA. JAK2 has been investigated the most in neutrophil function compared to the other JAK proteins. To further investigate the cellular mechanism by which JAK signaling influences the proteins associated with various neutrophil functions, we conducted mass spectrometry analysis, Luminex and functional assays. Prior to any analysis, we first verified that our treatments (PMA application and inhibitors) did not change the morphology of the cells (Supplemental Figure 1B). No differences were observed in cell diameter between the aged or young neutrophils in any of the conditions (Supplemental Figure 1C). Also, there were no observed sex differences in diameter in the young (Supplemental Figure 1D) or the age (Supplemental Figure 1E) neutrophils in any conditions.

### Sex and age directly influence neutrophil cellular profile with JAK2 inhibition

Mass spectrometry data was organized to determine uniquely expressed and shared proteins among the groups in different conditions and the shared proteins per each condition. The results revealed 2765 proteins were sampled from all groups in the secretome and 4273 in the cell fraction (Table 1). Unique proteins expression, driven by treatment condition, were also found in each compartment as detailed in Table 1. Interestingly, samples from the aged female neutrophils showed the highest number of uniquely expressed proteins in the secretome, as did neutrophils from young female in the cell fraction under all conditions (Table 1). Further characterization of the similarities across treatment conditions showed several shared proteins as detailed in Table 2. These data highlight the effect of both sex and age on JAK regulation of neutrophil cellular function (Figure 1).

**Figure 1:**
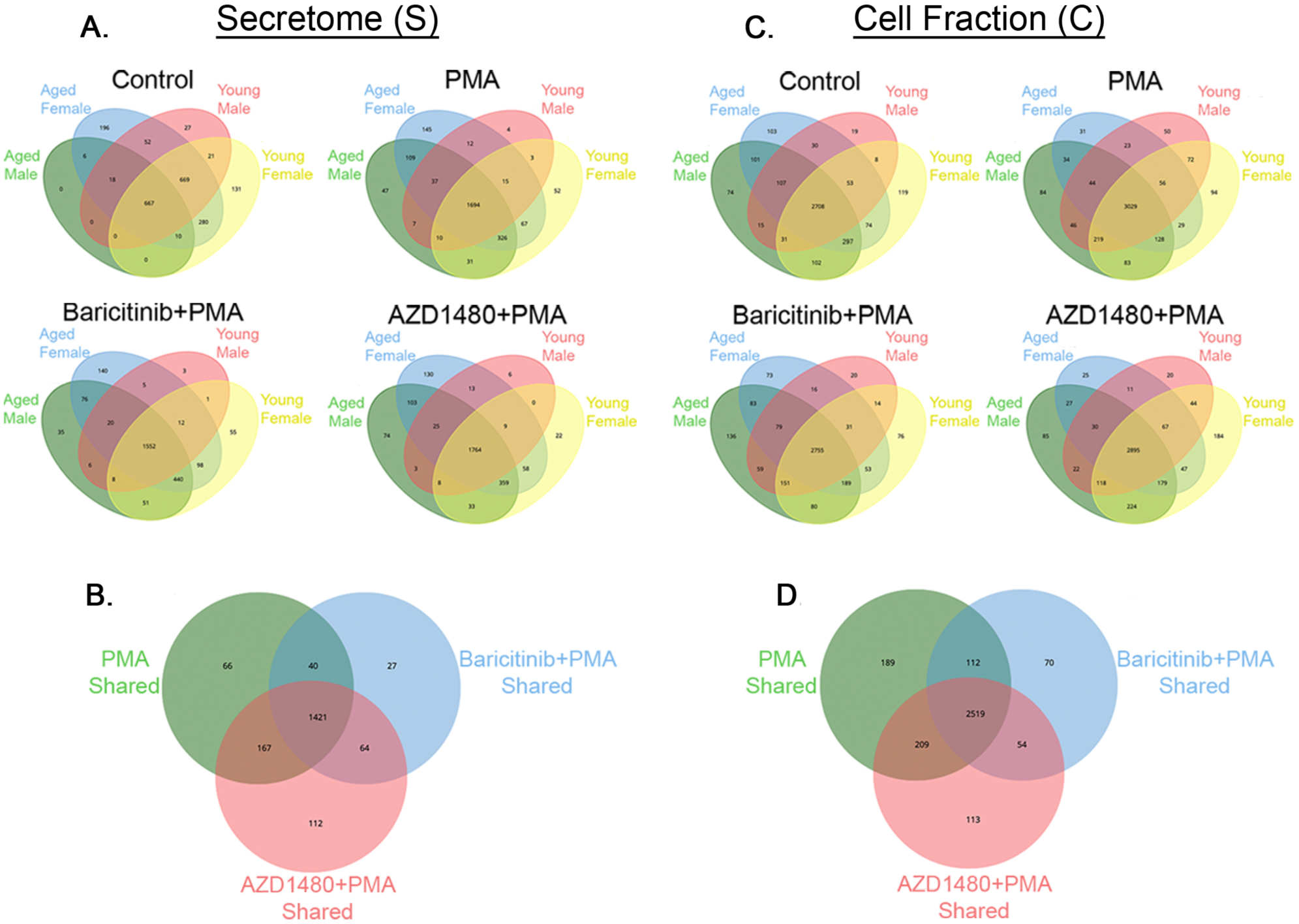
Age-, sex-, and condition-dependent changes in neutrophil dictate unique protein profiles, with shared protein networks regulating homeostasis. **A.** Venn diagrams representing unique and shared proteins identified in the secretome (S) across different experimental conditions (Control, PMA, Baricitinib+PMA, and AZD1480+PMA) and stratified by age and sex groups. **B.** Venn diagram representing all shared proteins from A in the secretome across conditions. **C.** Venn diagrams illustrating unique and shared proteins identified in the cell fraction (C) under treatment conditions. **D.** Venn diagram comparing shared proteins from C in the cell fraction across experimental conditions. YM= young male, YF= young female, AM= aged male, AF= aged female.

**Table 1:** There are different amounts of uniquely expressed proteins in each group.

|  |  |  | Uniquely Expressed Proteins |  |  |  |
| --- | --- | --- | --- | --- | --- | --- |
|  | Group: | Total proteins | Control | PMA | Baricitinib+PMA | AZD1480+PMA |
| **Secretome**<br>**(S)** | **YM** | 2765 | 27 | 4 | 3 | 4 |
|  | **YF** | 2765 | 13 | 52 | 55 | 52 |
|  | **AM** | 2765 | 0 | 47 | 35 | 74 |
|  | **AF** | 2765 | 196 | 145 | 140 | 130 |
| **Cell Fraction**<br>**(C)** | **YM** | 4273 | 19 | 50 | 20 | 20 |
|  | **YF** | 4273 | 119 | 94 | 76 | 184 |
|  | **AM** | 4273 | 74 | 84 | 136 | 85 |
|  | **AF** | 4273 | 103 | 31 | 13 | 25 |

**Table 2:**
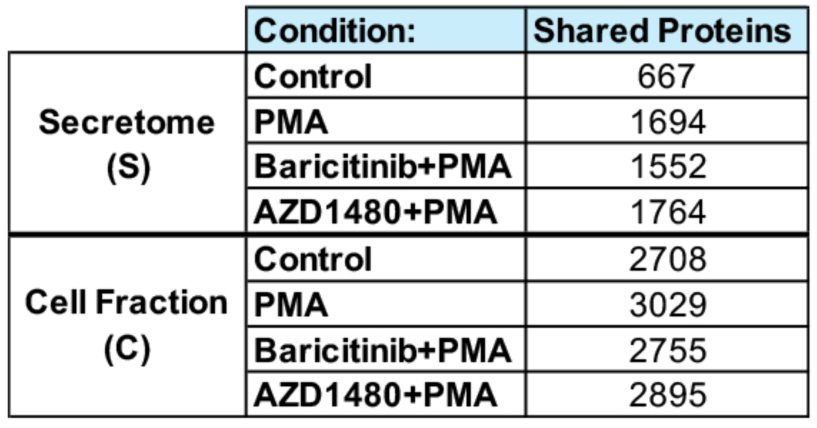
There are similar amounts of shared proteins in each condition.

#### Secretome

Under control conditions, we identified 27, 13, 0, and 196 uniquely expressed proteins in the young male, young female, aged male, and aged female groups, respectively, with 667 proteins shared among all groups (Figure 1A). The addition of PMA changed the number of uniquely expressed proteins (4, 52, 47, and 145), and increased the number of shared proteins across groups, 1694 detected proteins. Inhibition of all JAK proteins before PMA activation (Baricitinib + PMA) also altered the number of uniquely expressed proteins (3, 55, 35, and 140), with a slight reduction in the number of shared proteins (1552). Inhibition of JAK2 alone before PMA activation (AZD1480 + PMA) also increased the number of unique proteins, as compared to control (4, 52, 74, and 130), while increasing the number of shared proteins (1764) (Figure 1A). Direct comparisons of the shared proteins across treatment shows 1421 common proteins, with 66 proteins unique to PMA, 27 to Baricitinib + PMA, and 112 to AZD1480 + PMA (Figure 1B).

#### Cell Fraction

Under control conditions, we identified 19, 119, 74, and 103 uniquely expressed proteins in the young male, young female, aged male, and aged female groups, respectively, with 2708 proteins shared among all groups (Figure 1C). The addition of PMA changed the number of uniquely expressed proteins (50, 94, 84, and 31), and increased the number of shared proteins across groups, 3029 detected proteins. Inhibition of all JAK proteins before PMA activation (Baricitinib + PMA) altered the number of uniquely expressed proteins (20, 76, 136, and 13), with a reduction in the number of shared proteins (2755). Inhibition of JAK2 alone before PMA activation (AZD1480 + PMA) increased the number of unique proteins, as compared to control (20, 184, 85, and 25), while increasing the number of shared proteins (2895) compared to the Baricitinib + PMA group (Figure 1C). Direct comparisons of the shared proteins across treatment shows 2519 common proteins, with 189 proteins unique to PMA, 70 to Baricitinib + PMA, and 113 to AZD1480 + PMA (Figure 1D). Overall, all these results highlight both the shared and distinct proteins associated with neutrophil activation that are both dependent and independent of the JAK signaling pathway.

### JAK2 regulates proteins involved with neutrophil migration in an age-dependent manner

To determine how JAK activation contributes to neutrophil migration and whether age plays a role in this pathway, JAK signaling was inhibited in isolated neutrophils from young and aged mice, activated by PMA, and placed in a Boyden chamber (with 3 µm pores). Secretome was analyzed to determine the effect on cellular response to the inhibition, cell fraction to determine the effect of the inhibition on intracellular signaling, and functional migration assay to determine the effect of the inhibition on this neutrophil function. Cell migration is the result of physiological change associated with intracellular modification, including cytoskeletal remodeling to form pseudopodia (cell fraction and migration analysis), in response to various extracellular cues (secretome and cytokine analysis).

#### Secretome

The secretome analysis of our samples provide insights into how JAK signaling affects the release of extracellular factors by neutrophils, reflecting dynamic changes in protein secretion. Our data show that Insulin-like Growth Factor (IGF) pathway activity is regulated by JAK2, as seen by the increased proteins in this pathway under JAK2 inhibition (AZD1480) in young and aged neutrophils (Supplemental Figure 2A). IGF plays a crucial role in plasma membrane dynamics, as increased IGF signaling promotes lipid incorporation into the plasma membrane, leading to increased stiffness and reduced cellular motility [34]. 3β-hydroxysterol Δ24-reductase (DHCR24), a key regulator of cholesterol levels and plasma membrane stiffness [35], is upregulated upon JAK2 inhibition in young and aged neutrophils (Supplemental Figure 2A). Chemokine signaling, essential for directing neutrophil migration, also show JAK2-dependent modulation. In young neutrophils, JAK2 regulates specific chemokine signaling (CXCL2), seen by the increase in the proteins in this pathway under JAK2 inhibition (Supplemental Figure 2A) with no effect on chemokine release (Supplemental Figure 2B). In aged neutrophils, JAK2 increases release of CXCL2 (p=0.0279; Supplemental Figure 2B) with no effect on chemokine signaling (Supplemental Figure 2A). Additionally, the secretion of IL-1α, a pro-inflammatory cytokine that enhances neutrophil recruitment and activation, is upregulated in both young and aged neutrophils following JAK2 inhibition (Supplemental Figure 2B). The secretion of VEGF, a potent angiogenic factor, was also significantly increased in aged neutrophils following JAK2 inhibition (p<0.0001; Supplemental Figure 2B). Also, SDF1α secretion was increased in young neutrophils with JAK2 inhibition (p=0.0004; Supplemental Figure 2B). These findings suggest that JAK2 inhibition differentially regulates neutrophil secretion of key proteins in an age dependent manner.

Interestingly, our data show that the proteins modulated by JAK2 inhibition alone were also inhibited by pan-JAK inhibition, including IGF, DHCR24, IL-1α, SDF1α (young only) and VEGF. These data suggest that JAK2 may have the most direct effect on the regulation of these proteins. There were a few pathways that were unique to the pan-JAK modulation. SDF1α secretion was significantly increased in aged neutrophils following pan-JAK inhibition (p=0.0196; Supplemental Figure 2B). C1qR a complement receptor promoting neutrophil migration was significantly elevated in both young (p=0.0049) and aged (p<0.0001) neutrophils following pan-JAK inhibition (Supplemental Figure 2B).

#### Cell Fraction

Actin cytoskeleton rearrangement is crucial for neutrophil migration, and our data indicate that JAK2 signaling plays a significant role in regulating this process in an age-dependent manner. In young neutrophils, actin cytoskeleton signaling is influenced by JAK2 activation, as seen by the regulation of the ARP-WASP complex, a key driver of actin remodeling (Figure 2A) [36]. JAK2 inhibition increased Nuclear-factor-kappa-light-chain-enhancer of activated B cells 1 (NF*κ*B1), which regulates neutrophil migration through MAPK signaling [37], in aged neutrophils (Figure 2B). I-kappa-B kinase 2, responsible for phosphorylating MAPK, decreased with JAK2 inhibition in aged but not young neutrophils (Figure 2B). Additionally, Actin-Related Protein 2/3 Complex (ARP2/3), which is essential for cytoskeletal rearrangement [38], is decreased by JAK2 inhibition in young neutrophils, ARPC2 (p=0.0125; Figure 2C) and ARPC3 (p=0.022; Figure 2C). In aged neutrophils, JAK2 inhibition also decreased ARPC2 levels (p=0.018; Figure 2C), and ARPC3 levels (p=0.0474; Figure 2C). Myristoylated Alanine-Rich Protein Kinase C (MARCKS) pathway, a critical regulator of ARP2/3 phosphorylation and actin nucleation [39, 40], is upregulated by JAK2 inhibition in both young and aged neutrophils (Figure 2B). However, JAK2 had no effect on MARCKS protein levels in either young or aged neutrophils (Figure 2C). These findings suggest that JAK2 signaling predominantly regulates migration through cytoskeletal remodeling in both young and aged neutrophils.

**Figure 2:**
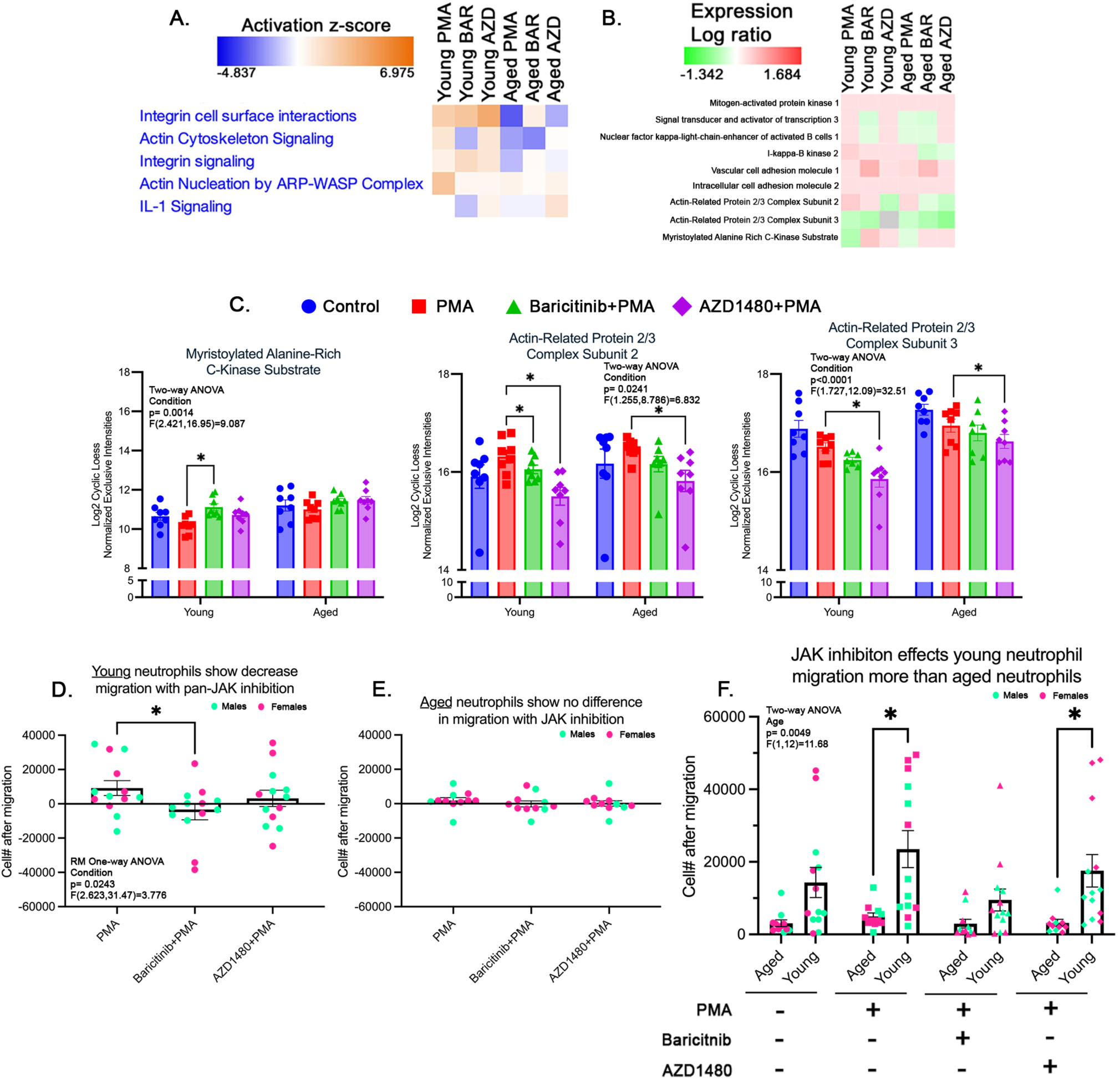
There are age-dependent changes in JAK2 regulation of neutrophil migration. **A.** Functional heatmap of cell fraction depicting the activation z-scores of migration-associated pathways (normalized to control). **B.** Heatmap of protein expression changes (log ratio) in proteins associated with neutrophil motility (normalized to control). **C.** Bar graphs showing the relative expression of: MARCKS, young Bar is significantly increased compared to PMA. ARPC2, young and aged AZD are significantly decreased compared to PMA, young Bar is significantly decreased compared to PMA. ARPC3, young and aged AZD are significantly decreased compared to PMA (n=8/condition/age). **D-E.** Boyden chamber quantification of neutrophil migration (normalized to control), stratified by age, showing a decrease in young neutrophil migration under pan-JAK inhibition, no difference in migration of aged (young: n=13/condition; aged: n=11/condition). **F.** Comparison of raw migration data shows significant increase in neutrophil migration of young neutrophils compared to aged under PMA and AZD (young: n=13/condition; aged: n=11/condition). p-value: * <0.05; bars represent mean ± SEM. Bar= Barcitinib treated, AZD= AZD1480 treated.

Pan-JAK inhibition showed an interesting regulatory pattern in terms of modulating neutrophil migration. In aged neutrophils, integrin cell surface interactions were upregulated by pan-JAK inhibition (Figure 2A). IL-1 signaling, known to influence neutrophil migration via CXCR2 [41], was decreased by the inhibition of all JAK proteins in young neutrophils, with no effect in aged neutrophils (Figure 2A). NF*κ*B1 was decreased when all JAK proteins were inhibited in young neutrophils (Figure 2B). Additionally, I-kappa-B kinase 2, was decreased with pan-JAK inhibition in aged neutrophils (Figure 2B). MARCKS protein level was increased in young neutrophils with pan-JAK inhibition (p=0.0224; Figure 2C).

Functional assessment of neutrophil migration was done using the Boyden chamber with the chemoattractant IL-8. In agreement with the data observed in the secretome and cell fraction mass spectrometry analyses, the inhibition of JAK led to fewer migrating cells in young neutrophils (p=0.036; Figure 2D). JAK activity had no effect on the migration of aged neutrophils (Figure 2E). Combined with the mass spectrometry and Luminex data, these data support the age dependent effect of JAK modulation of neutrophil migration. A direct comparison of the young and aged neutrophils showed more migration in young neutrophils (pan-JAK: p=0.0191; JAK2: p=0.0169, Figure 2F). Specifically, pan-JAK inhibition decreased migration of young neutrophils compared to aged neutrophils (Supplemental Figure 3A). These age specific changes were driven by increased migration of young female neutrophils (Supplemental Figure 3B). No sex differences were observed in JAK regulation of neutrophil migration within age groups (Supplemental Figure 3C and 3D).

**Figure 3:**
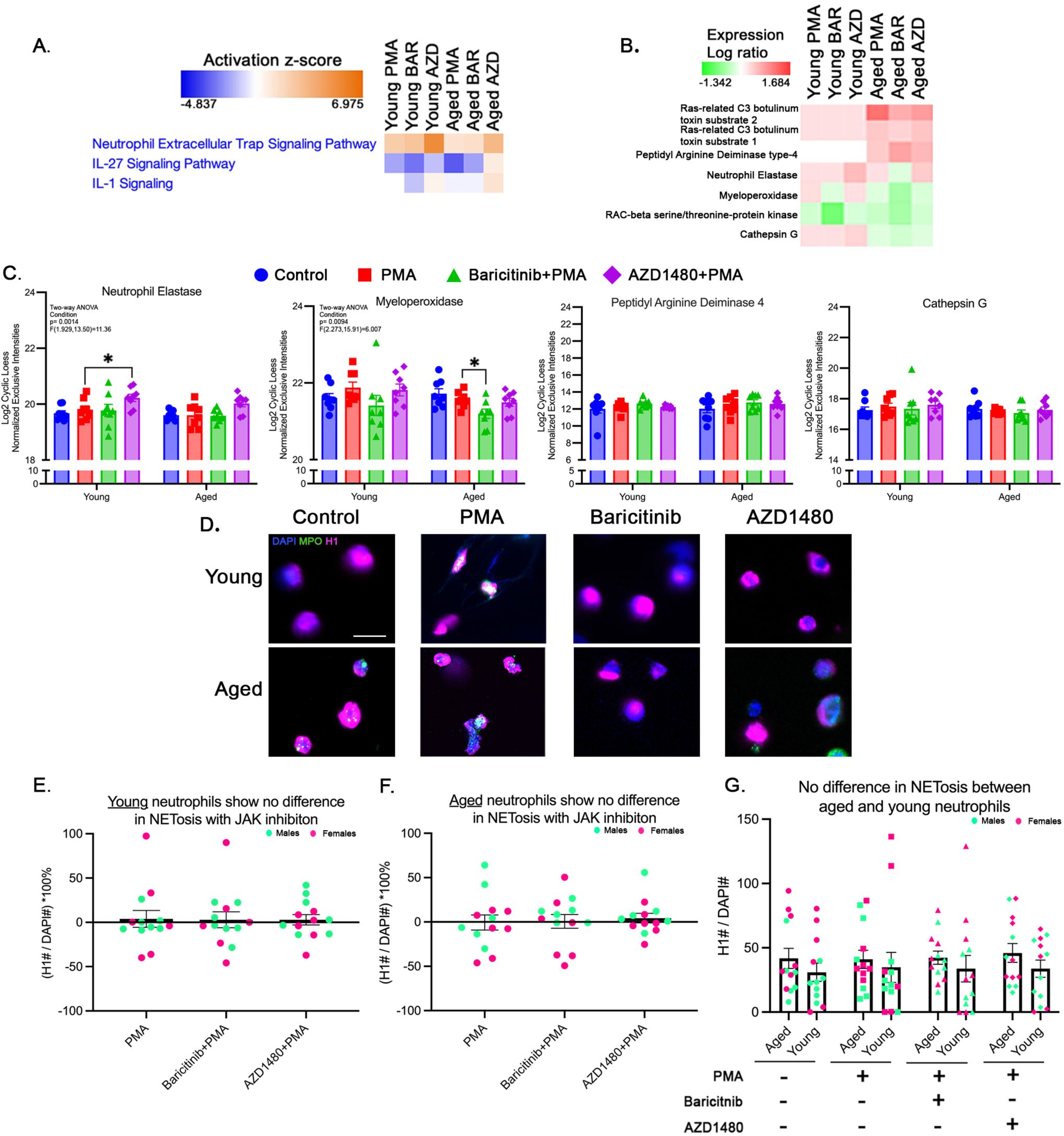
JAK2 does not regulate NETosis in young or aged neutrophils. **A.** Functional heatmap of cell fraction displaying activation z-scores for pathways associated with NETosis (normalized to control). **B.** Heatmap illustrating protein expression (log ratio) changes in NETosis-related proteins (normalized to control). **C.** Bar graphs showing the relative expression of NETosis-associated proteins, NE, MPO, PAD4, and cathepsin G, across experimental conditions (n=8/condition/age). **D.** Representative images of NET formation in young and aged neutrophils under all conditions by DNA (DAPI), histone H1, and MPO staining. **E-F.** Quantification of NETosis in young and aged neutrophils (normalized to control), showing no significant differences with JAK inhibition (young: n=13/condition; aged: n=12/condition). **G.** Raw comparison of the number of NETosing cells between young and aged neutrophils (young: n=13/condition; aged: n=12/condition). p-value: * <0.05; bars represent mean ± SEM; scale bar= 20µm

### JAK2 activation does not contribute to NET production, despite its modulation of neutrophil elastase levels

Neutrophil extracellular traps (NETs) are an important effector function of neutrophils. NETosis, the molecular mechanism of NET release, is characterized by the release of decondensed DNA speckled with granule proteins. NETosis is dependent on ROS production, MPO activity, NE activity, and the activation of peptidylarginine deaminase 4 (PAD4) [42]. Cathepsin G induces PAD4-dependent NETosis by facilitating chromatin decondensation and NET formation [43]. PMA induces ROS-dependent lytic NETosis via the PKC/Raf/MEK/ERK pathway [44]. Here we investigated if JAK signaling is required for NETosis.

#### Secretome

The secretome analysis revealed significant JAK2-dependent changes in NET-associated protein regulation, with distinct age-related effects. In young neutrophils, JAK2 inhibition increased NET signaling pathway activity, whereas in aged neutrophils, JAK2 inhibition led to a downregulation of NET signaling (Supplemental Figure 4A). Extracellular localization of Matrix Metalloproteinase-8 (MMP-8), a granule enzyme critical for extracellular matrix remodeling, and S100 calcium-binding protein A8 (S100A8), a cytoplasmic calcium-sensing protein, are both associated with NET release [45, 46]. Luminex analysis revealed that JAK2 inhibition increased MMP-8 only in aged neutrophils, and S100A8 secretion in both young and aged neutrophils (Supplemental Figure 4B).

Just like with JAK2 only inhibition, pan-JAK inhibition led to an increase in young neutrophils, and a decrease in aged neutrophils (Supplemental Figure 4A) of proteins associated with NET signaling pathway. Both MMP8 and S100A8 levels were increased in the extracellular space of both young and aged neutrophil cultures (Supplemental Figure 4B).

#### Cell Fraction

Our analysis of the NET signaling pathway revealed distinct JAK2-dependent regulatory effects in young and aged neutrophils. In both age groups, JAK2 inhibition led to an increase in NET signaling (Figure 3A). Previous studies indicate that IL-27 plays a role in NET formation through the JAK1/2-STAT1/3 pathway in human peripheral neutrophils [47]. Our data show that IL-27 signaling is modulated by JAK2 inhibition exclusively in aged neutrophils (Figure 3A). Additionally, IL-1β signaling, which has been linked to NETosis [48, 49], was differentially affected by JAK2 inhibition, leading to an increase in both young and aged neutrophils (Figure 3A). Mass spectrometry analysis further revealed that JAK2 regulates key NET-associated proteins, including NE and MPO. JAK2 inhibition significantly increased NE activity in young neutrophils (Figure 3B), a result that corresponded with an increase in NE protein levels in AZD1480-treated young neutrophils (p=0.0134; Figure 3C). MPO levels remained unchanged with JAK2 inhibition in both young and aged neutrophils (Figure 3B). Further characterization of PAD4 and cathepsin G, both critical regulators of NETosis, showed age-dependent activity but no direct regulation by JAK2 (Figure 3B and 3C).

In contrast to JAK2 inhibition, pan-JAK inhibition reduced IL-1β signaling in both young and aged neutrophils (Figure 3A), suggesting a regulatory role for other JAK family members in modulating NET-associated inflammation. MPO levels significantly decreased following pan-JAK inhibition in young and aged neutrophils (Figure 3B), with further analysis showing a specific reduction in MPO protein levels in aged neutrophils (p=0.0238; Figure 3C). Unlike JAK2 inhibition, which selectively influenced NE activity, pan-JAK inhibition did not alter NE protein levels in either age group (Figure 3C). Similar to JAK2 inhibition, PAD4 and cathepsin G displayed age-dependent activity but were unaffected by pan-JAK inhibition (Figure 3B and 3C). These findings suggest that each JAK family protein plays a distinct modulatory role in NETosis, with JAK2 playing a direct role in regulating NE activity.

Functional analysis of NET production was conducted as described. Using Histone1 (H1) [50], myeloperoxidase (MPO) and DNA (DAPI) staining, we quantified the number of cells producing NETs in culture under all conditions. Representative images (Figure 3D) show the presence of NETs in all conditions. Our data show that NET production is not affected by JAK inhibition in both young and aged neutrophils (Figure 3E-G). No age or sex differences were observed under any of the conditions presented (Supplemental Figure 5). Collectively, these findings indicate that neutrophils from both young and aged mice produce NETs independently of JAK activation.

### JAK2 regulates proteins involved in ROS production in both young and aged neutrophils but modulates ROS-associated degranulation in an age-dependent manner

#### ROS Production

ROS production in neutrophils is primarily mediated by NADPH oxidase (NOX). NADPH oxidase complex is formed by the activation and translocation of p47phox, p40phox, and p67phox proteins to the plasma membrane [51], where they bind to the membrane-bound subunits gp91phox and p22phox, forming the active flavocytochrome b558 subunit. The formation of this subunit recruits Ras-related C3 botulinum toxin substrate 1/2 (RAC1/2) to the subunit forming a fully activated NADPH oxidase complex. Several studies have demonstrated that the activation of NADPH oxidase is influenced by several serine-threonine kinases, including Protein Kinase C (PKC) and MAPK [52]. Previous research shows contradictory role of JAK signaling in ROS production by human neutrophils. In rheumatoid arthritis, JAK inhibition had no effect on ROS production [53]. In asthma and COPD, inhibition of JAK signaling led to decreased ROS production [54]. These suggest that JAK modulation of neutrophil activation may be context dependent. In our study, we will determine whether JAK activation is critical to ROS production outside of disease context.

#### Secretome

The PI3K/AKT pathway regulates ROS production through both NOX activation and mitochondrial modulation. Upon activation, PI3K generates PIP3, leading to AKT activation, which enhances NOX complex assembly and subsequent ROS generation [55]. Our data demonstrates that JAK2 inhibition increases PI3K/AKT signaling in young neutrophils (Supplemental Figure 6). Whereas, in aged neutrophils, JAK2 inhibition decreases the PI3K/AKT pathway. ROS and NOS production signaling pathways are increased under JAK2 inhibition in the young neutrophils (Supplemental Figure 6). Conversely, in aged neutrophils, this pathway is decreased with JAK2 inhibition. Both the PI3K/AKT and the ROS and NOS production pathways were increased with pan-JAK inhibition in both young and aged neutrophils (Supplemental Figure 6).

#### Cell fraction

Our analysis of neutrophil cytoplasmic content revealed JAK2-dependent regulation of several pathways essential for ROS production. The SCF-KIT signaling pathway, known to regulate mitochondrial function and promote an oxidative neutrophil phenotype during cancer [56], was increased by JAK2 inhibition in both young and aged neutrophils (Figure 4A). The fMLP pathway, a well-established activator of ROS production [57], was also differentially modulated by JAK2. In young neutrophils, proteins associated with fMLP signaling decreased following JAK2 inhibition, while in aged neutrophils, JAK2 inhibition led to an increase in this pathway (Figure 4A). The PI3K/AKT pathway, which facilitates NOX1 activation via phosphorylation of p47phox, was not affected by JAK2 inhibition in young or aged neutrophils (Figure 4A). Furthermore, IL-1 signaling, a key driver of neutrophil ROS production [58], increased following JAK2 inhibition in young and aged neutrophils (Figure 4A). In young neutrophils NOX1, a critical component of ROS production, is decreased with JAK2 inhibition (Figure 4B). Additionally, levels of NOX1 were decreased with JAK2 inhibition in both young (p=0.002) and aged (p=0.0253) neutrophils (Figure 4C). Additionally, in aged neutrophils, JAK2 inhibition selectively downregulated ROCK2, a key activator of the NOX complex (Figure 4B).

**Figure 4:**
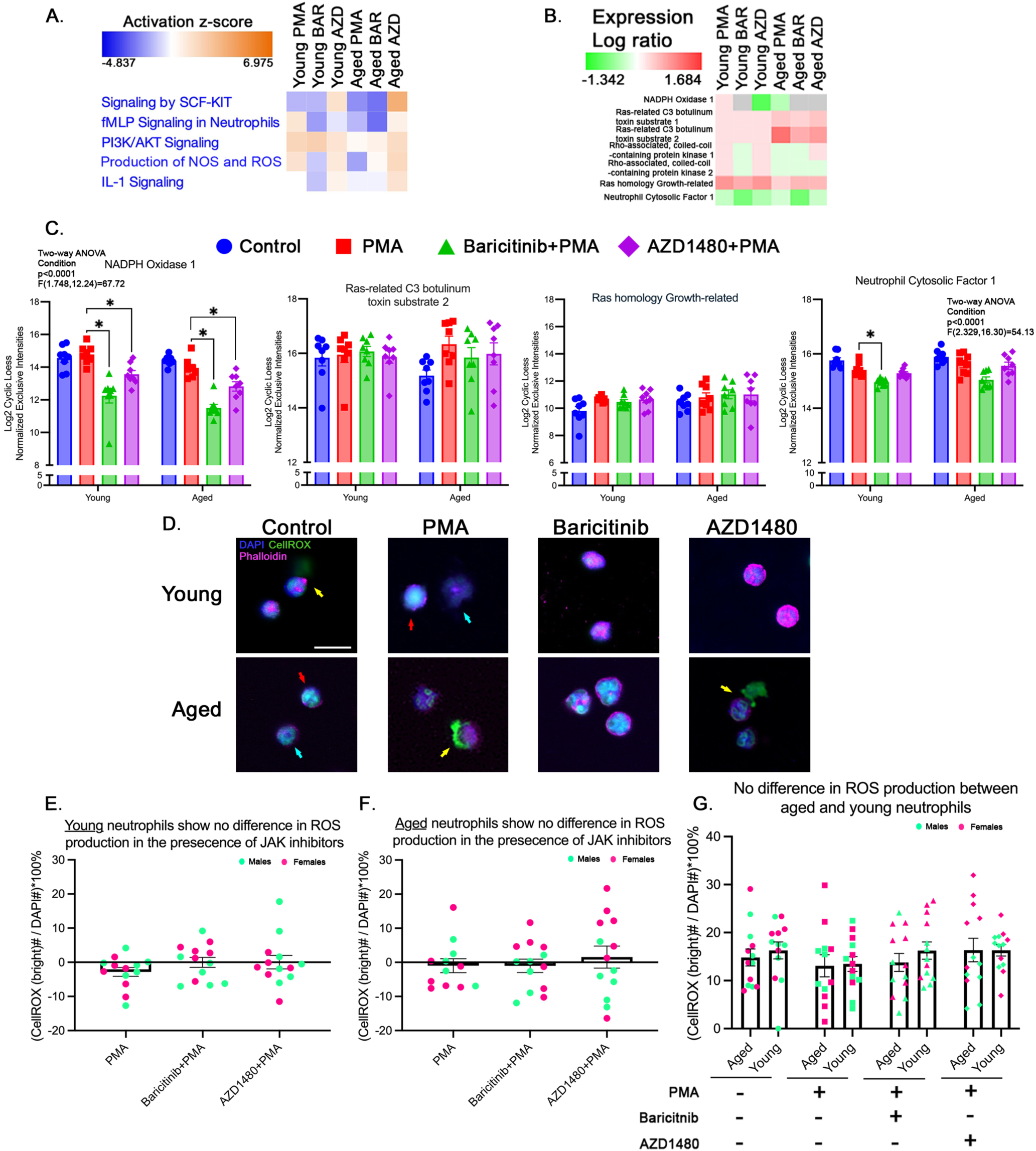
JAK2 signaling regulates key proteins involved in ROS production; however, it does not enhance ROS production. **A.** Functional Heatmap of cell fraction activation z-scores for ROS production-associated pathways (normalized to control). **B.** Heatmap depicting changes in protein expression (log ratio) levels involved in ROS generation (normalized to control). **C.** Bar graphs showing the relative expression of key proteins in ROS production, NOX1, RAC2, RhoG, and NCF1, across experimental conditions (n=8/condition/age). **D.** Representative images in young and aged neutrophils under all conditions stained with DAPI, CellROX green, and phalloidin to assess ROS production. **E-F** Quantification of ROS production in young and aged neutrophils (normalize to control) demonstrating no significant changes with JAK inhibition (young: n=13/condition; aged: n=12/condition). **G.** Raw comparison of cell’s intracellular ROS production between young and aged neutrophils showing no significant differences (young: n=13/condition; aged: n=12/condition). scale bar= 20µm; red arrow= increased ROS production, blue arrow= no change in ROS production, yellow arrow= ROS-associated degranulation; bars represent mean ± SEM; p-value: * <0.05.

In young and aged neutrophils, fMLP signaling proteins were reduced following pan-JAK inhibition (Figure 4A). Whereas the ROS and NOS production pathway was decreased in the young neutrophils and increased in the aged neutrophils, following pan-JAK inhibition (Figure 4A). IL-1 signaling was suppressed by pan-JAK inhibition in young neutrophils (Figure 4A). NOX1 protein levels were significantly reduced in both young (p=0.0027) and aged (p=0.0002) neutrophils following pan-JAK inhibition (Figure 4C). Unlike JAK2 inhibition, which selectively affected ROCK2 in aged neutrophils, pan-JAK inhibition led to a reduction in both ROCK1 and ROCK2 in young and aged neutrophils (Figure 4B). Additionally, Neutrophil Cytosolic Factor 1 (NCF1), a key NOX1/2 complex subunit, remained unaffected by JAK2 inhibition but was significantly downregulated in young neutrophils with pan-JAK inhibition (p=0.0152; Figure 4C).

These findings suggest that the mechanisms underlying the modulatory effect of each JAK family protein on neutrophil ROS production may be unique.

The functional analysis of ROS production was determined using CellROX green in cultured neutrophils (Figure 4D; red arrow). JAK inhibition had no direct effect in neutrophil upregulation of ROS production across age (Figure 4E-G) or sex (Supplemental Figure 7C and D). Overall, the proteomic and functional analyses suggest that although JAK activity may influence the levels of key proteins in ROS production, in an age dependent manner, it does not lead to the production of ROS by neutrophils.

### ROS Degranulation

Degranulation in neutrophils is the process by which granules containing antimicrobial proteins, enzymes, and ROS are released into the extracellular space to combat pathogens. This tightly regulated process plays a pivotal role in innate immunity, balancing effective microbial killing with the prevention of excessive tissue damage [59]. NOX is a protein complex associated with primary neutrophil granules [60]. Neutrophil degranulation is a process often associated with the release of tertiary (gelatinase), secondary (specific), and primary (azurophilic) granules. With primary granules containing potent enzymes like MPO and NE, the threshold of release is higher due to their toxic effects [61]. Secondary granules store lactotransferrin (LTF) and lipocalin-2 (LCN2) to limit bacterial growth and tertiary granules contain metalloproteases. The fusion of neutrophil granules with the plasma membrane or phagosomes is mediated by SNARE proteins, including syntaxins, VAMPs, and SNAPs, which coordinate vesicle docking and membrane fusion for efficient secretion of antimicrobial contents [62]. Calcium signaling plays a crucial role in neutrophil degranulation by activating actin cytoskeletal remodeling and triggering granule transport through motor proteins [63].

#### Secretome

Our data indicate that JAK2 modulation of fMLP signaling in neutrophils is age dependent. In aged neutrophils, JAK2 inhibition decreased the activity of this pathway (Figure 5A). RAC signaling, which is crucial for neutrophil degranulation through the reorganization of the actin cytoskeleton, is also influenced by JAK2 in an age-dependent manner. Inhibition of RAC activity disrupts granule fusion and attenuates the release of antimicrobial mediators [64]. Our data show that JAK2 inhibition leads to a decrease in RAC signaling in aged neutrophils, with no effect in young neutrophils (Figure 5A). Characterization of tertiary granule proteins release revealed that MMP-8 is increased by JAK2 inhibition only in aged neutrophils (Supplemental Figure 4B), with no effect on MMP-9 levels (Supplemental Figure 8). Secondary granule proteins, such as LTF and LCN2, were both increased with JAK2 inhibition in young and aged neutrophils (Figure 5B). Upon further analysis, in aged neutrophils, both LTF (p<0.0001) and LCN2 (p=0.0001) were significantly increased compared to PMA condition (Figure 5C). In young neutrophils only LTF was increased with JAK2 inhibition (p=0.0023) compared to PMA condition (Figure 5C). Release (extracellular) of primary granule proteins, including MPO and NE, were not affected by JAK2 inhibition in both young and aged neutrophils (Figure 5B). However, upon further analysis there was a significant decrease in NE protein levels with JAK2 inhibition in aged neutrophils (Figure 5C). Cytokine release, associated with neutrophil secretory vesicles, was also regulated in an age dependent manner. The anti-inflammatory cytokine IL-10 increased (p=0.0413) under JAK2 inhibition in the young neutrophils (Supplemental Figure 8).

**Figure 5:**
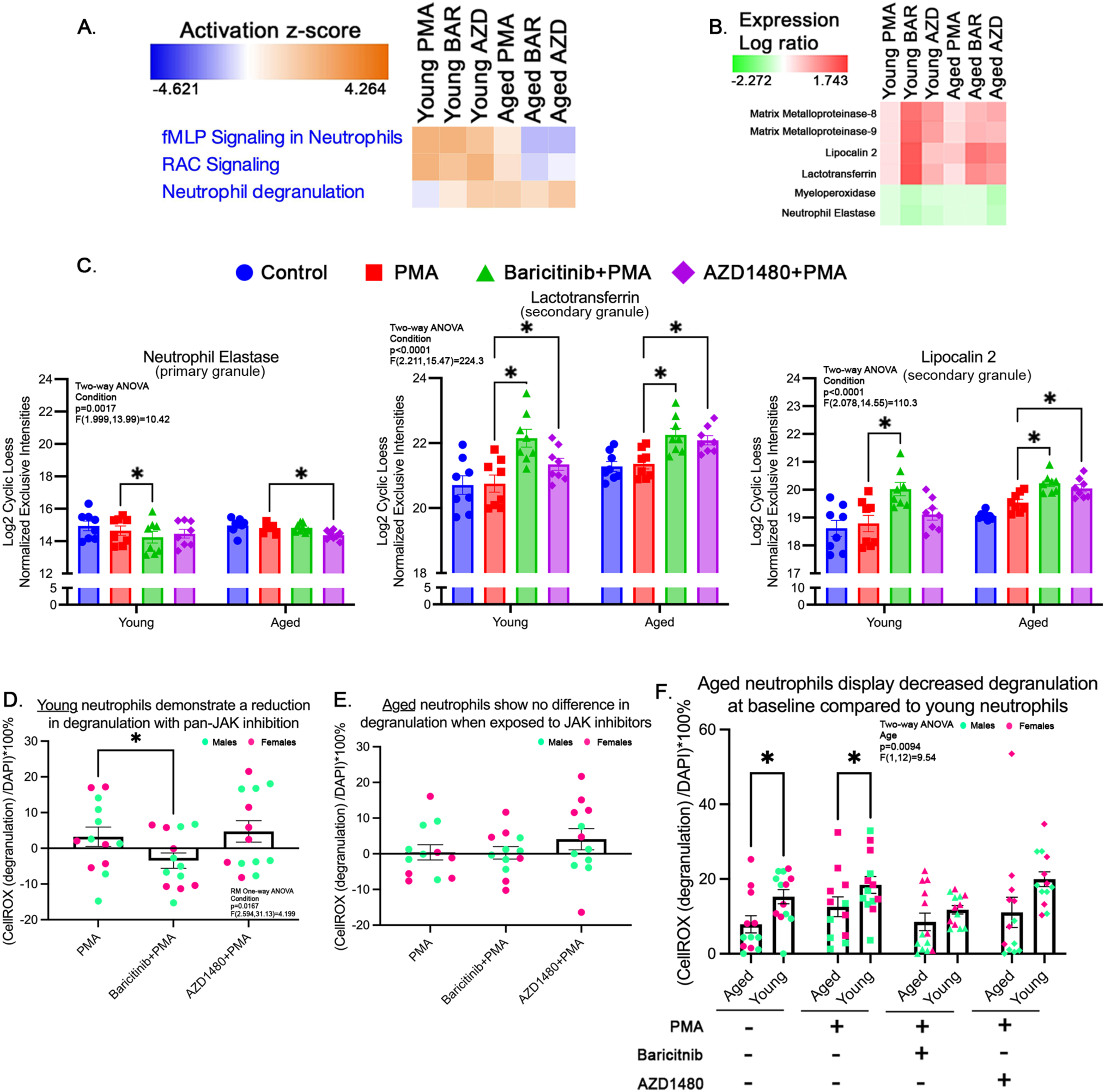
JAK2 signaling regulates ROS release (proxy for degranulation) in young neutrophils, but not in aged neutrophils. **A.** Functional heatmap of secretome showing activation z-scores for pathways related to degranulation (normalized to control). **B.** Heatmap depicting protein expression (log ratio) changes in degranulation-associated pathways (normalized to control). **C.** Bar graphs displaying expression levels of granule-associated proteins, NE (primary granule protein), LTF (secondary granule protein), and LCN2 (secondary granule protein) across experimental conditions (n=8/condition/age). **D-E.** Quantification of degranulating (ROS releasing) neutrophils in young and aged neutrophils (normalized to control) under different conditions shows a reduction with pan-JAK inhibition in young neutrophils (n=13/condition; aged: n=12/condition). **F.** Raw comparison of degranulation between young and aged neutrophils demonstrates decreased degranulation under control and PMA treatment in aged neutrophils (n=13/condition; aged: n=12/condition). p-value: * <0.05; bars represent mean ± SEM; NE: neutrophil elastase; LTF: lactotransferrin; LCN2: lipocalin2.

Interestingly, pan-JAK inhibition had the same effect on fMLP and RAC signaling pathways as JAK2, suggesting that JAK2 may be driving the changes observed in these pathways (Figure 5A). Investigating the tertiary granule release, pan-JAK inhibition increased MMP-8 secretion in both young (p=0.0094) and aged (p=0.0005) neutrophils (Supplemental Figure 4B), with no effect on MMP-9 (Supplemental Figure 8). Secondary granule proteins, including LTF and LCN2, showed an increase compared to PMA condition in both young and aged neutrophils (Figures 5B and 5C). Additionally, primary granules release, such as NE and MPO, were not affected in young or aged neutrophils with pan-JAK inhibition (Figure 5B and 5C). However, upon further analysis, there was a significant decrease in NE in young neutrophils with pan-JAK inhibition (Figure 5C). Analysis of cytokine secretion revealed that pan-JAK inhibition significantly increased secretion of TNF-α (p=0.0025) and CXCL10 (p=0.0114) in aged neutrophils (Supplemental Figure 8). IL-10 (p=0.0089) and CXCL10 (p=0.0001) levels were elevated in young neutrophils under pan-JAK inhibition (Supplemental Figure 8). The secretion of Serpin E1 and MMP-9 was not affected by JAK inhibition in either age group (Supplemental Figure 8).

Functional characterization of degranulation was performed by characterizing the release of ROS in the extracellular space, cells with adjacent CellRox signal outside the phalloidin stain (Figure 4D; yellow arrow) as a proxy for degranulation. Only pan-JAK inhibition significantly reduced ROS degranulation in young neutrophils (p=0.0344; Figure 5D). JAK inhibition had no effect on degranulation in aged neutrophils (Figure 5E). A few age differences were observed in ROS degranulation between young and aged neutrophils at baseline (p=0.028) and with PMA activation (p=0.0295; Figure 5F). Of note, these differences were abolished with JAK inhibition (Figure 5F). Correlation analysis shows that ROS degranulation is more affected in young neutrophils compared to aged neutrophils with the loss of all JAK activity (Supplemental Figure 10A), which seems to be driven by yet unknown differences between young and aged male neutrophils (Supplemental Figure 10B). No sex differences were observed in degranulation in young neutrophils (Supplemental Figure 10C and D). However, aged male neutrophils show a decrease of ROS-associated degranulation compared to aged female neutrophils with PMA activation (p=0.0386) and pan-JAK inhibition (p=0.0185; Supplemental Figure 10D).

#### Cell fraction

Our data indicates that JAK2 modulation of neutrophil degranulation is age dependent. Granulocyte Macrophage-Colony Stimulating Factor (GM-CSF) signaling, known to enhance degranulation through activation of PI3K/AKT and MAPK [65, 66], is upregulated in both young and aged neutrophils following JAK2 inhibition (Supplemental Figure 9). RAC signaling is decreased in young and aged neutrophils with JAK2 inhibition (Supplemental Figure 9). Intracellular calcium signaling was elevated in both young and aged neutrophils following JAK2 inhibition (Supplemental Figure 9). Similarly, SNARE signaling, which regulates the fusion of granules to the plasma membrane, is upregulated across both age groups (Supplemental Figure 9).

In agreement with the functional assessment, pan-JAK inhibition decreased the activity of several proteins involved in neutrophil degranulation. In young neutrophils, pan-JAK inhibition reduces GM-CSF, calcium, RAC and SNARE signaling (Supplemental Figure 9). In aged neutrophils, pan-JAK inhibition only decreased GM-CSF and RAC signaling pathways (Supplemental Figure 9). Our data suggest that RAC and SNARE signaling are critical to degranulation, as previously demonstrated [62, 64]. In addition, this suggests that overall, the effect of pan-JAK activation on neutrophil degranulation wanes with age. Taken together, these findings indicate that while JAK2 selectively regulates key components of degranulation it is not sufficient to trigger degranulation in neutrophils.

### JAK2 regulates the shift of energy metabolism with age

All the above listed neutrophil functions require energy, so the next cellular process we investigated was glycolytic metabolism. Neutrophils primarily use glycolysis for energy generation [67]. Neutrophil activation using PMA directly impacts metabolism by increasing oxygen consumption and activation of the pentose phosphate pathway (PPP) [68]. Whether this is influenced by JAK signaling is yet unknown. In this study, we assessed the effect of JAK inhibitors on neutrophil glycolytic output. Glycolytic metabolism was assessed in neutrophils using the Seahorse XF analyzer (XF96).

#### Secretome

Our data indicate that JAK2 inhibition decreases glucose and glycogen metabolism in young neutrophils (Supplemental Figure 11). Pan-JAK inhibition reduced glucose metabolism in both young and aged neutrophils (Supplemental Figure 11). Young neutrophils exhibit a decrease in pathways associated with the integration of energy metabolism, which facilitates rapid metabolic switching between glycolysis and mitochondrial respiration to sustain ATP production [69]. Furthermore, the PPP, essential for generating NADPH [70], is decreased in young neutrophils following pan-JAK inhibition (Supplemental Figure 11). Glycogen metabolism is decreased under pan-JAK inhibition in aged neutrophils (Supplemental Figure 11).

#### Cell fraction

Our data indicate that JAK2 modulation of glucose metabolism is decreased in both young and age neutrophils with JAK2 inhibition (Figure 6A). Glucose-6-phosphate dehydrogenase (G6PD), a key enzyme in glycolysis, is significantly reduced (p=0.0415) in young neutrophils following JAK2 inhibition (Figures 6B and 6C). JAK2 inhibition increased glycogen metabolism in aged neutrophils compared to the PMA condition (Figure 6A). Indeed, the activity of Glycogen debranching enzyme (AGL), which facilitates glycogen breakdown, is increased in aged neutrophils under JAK2 inhibition (Figure 6B). The PPP is also decreased by JAK2 inhibition in both young and aged neutrophils (Figure 6A). Ribose-5-Phosphate Isomerase, a PPP enzyme involved in the conversion of G6P to ribulose 5-phosphate [71], decreased in young neutrophils following JAK2 inhibition (Figure 6B). These findings suggest that JAK2 specifically regulates glycogen metabolism and PPP activity.

**Figure 6:**
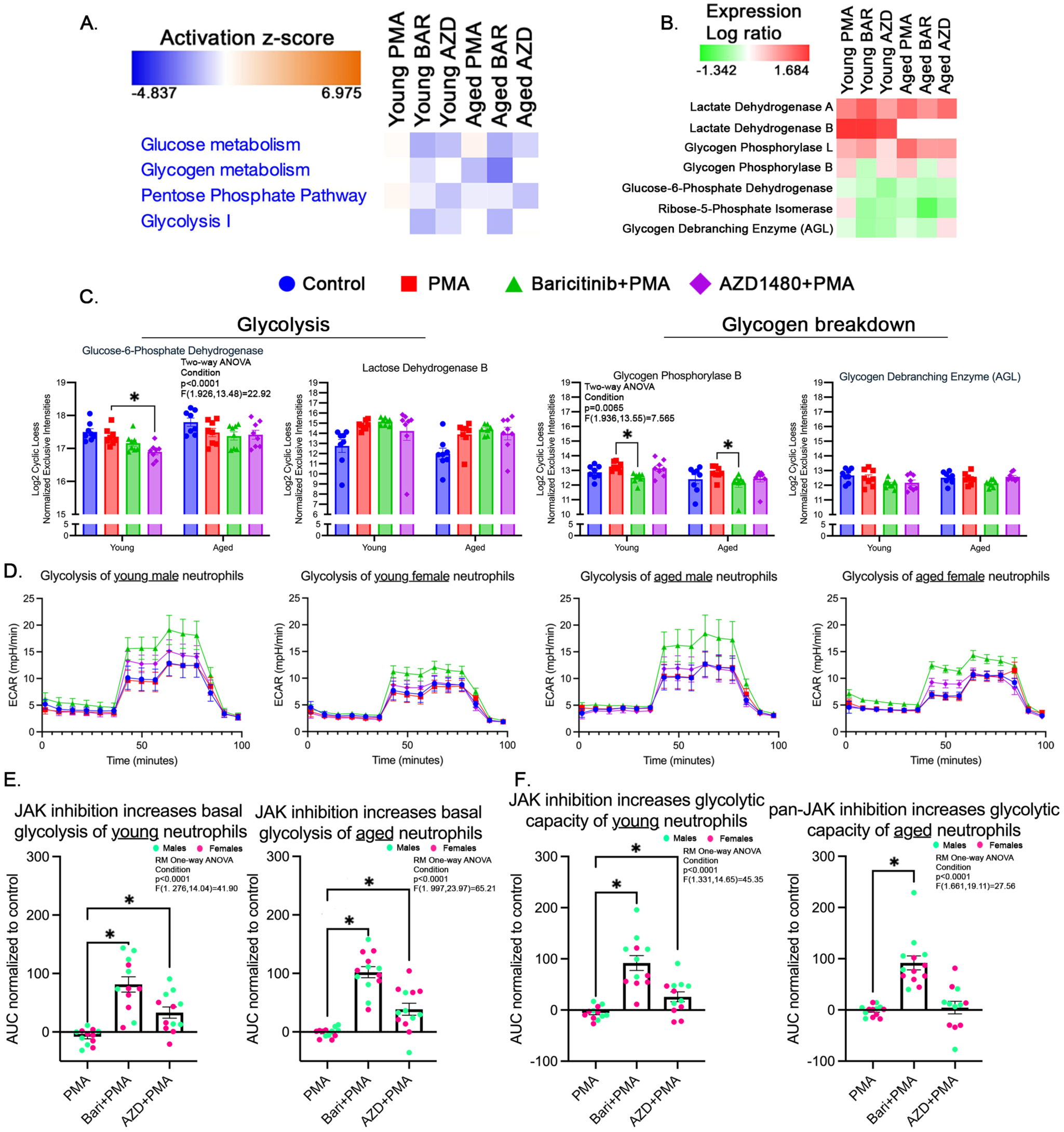
JAK2 inhibition increases basal glycolysis and glycolytic capacity in young and aged neutrophils. **A.** Functional heatmap illustrating activation z-scores for metabolic pathways (normalized to control). **B.** Heatmap of protein expression (normalized to control) changes in metabolic regulators. **C.** Bar graphs showing expression levels (log ratio) of key metabolic enzymes, glucose-6-phosphate dehydrogenase, Lactate dehydrogenase B, glycogen phosphorylase B, and glycogen debranching enzyme (AGL; n=8/condition/age). **D.** Representative histogram of Seahorse assay assessing glycolysis in neutrophils. ECAR is the acidification rate, i.e. lactate production, of cells under the various conditions. **E-F.** Quantification of ECAR using Area Under the Curve (AUC) of basal glycolysis and glycolytic capacity (normalized to control) in young and aged neutrophils. The data show increased glycolysis with JAK inhibition, in both age groups (n=12/condition; aged: n=12/condition). p-value: * <0.05; bars represent mean ± SEM.

Young neutrophils have decreased glucose metabolism, glycogen metabolism, PPP, and glycolysis following pan-JAK inhibition (Figure 6A). Aged neutrophils have decreased glucose metabolism and glycolysis with pan-JAK inhibition (Figure 6A). Lactate dehydrogenase A and B levels, which are critical for converting pyruvate to lactate in glycolysis, remain unaffected by JAK inhibition (Figure 6B and C). However, an age-related effect is observed on lactate dehydrogenase B levels in neutrophils, with aged neutrophils showing no change from control levels under all conditions (Figure 6B). Furthermore, the inactive “B” form of Glycogen Phosphorylase (PYGL), which converts glycogen into glucose-1-phosphate, is significantly downregulated in both young (p=0.0049) and aged (p=0.0217) neutrophils following pan-JAK inhibition, whereas the active “L” form remains unchanged (Figures 6B and 6C).

Functional characterization of the effect of JAK inhibition on neutrophils metabolism was performed using Seahorse metabolic assessment (Figure 6D). We analyzed the basal glycolysis, maximal glycolytic (capacity), and the trend between these two parameters in young and aged neutrophils. Except for glycolytic capacity in the aged neutrophils with JAK2 inhibition, pan-JAK and JAK2-only inhibition led to an increase in both basal and glycolytic capacity in all conditions (Figure 6E and 6F). Our data shows no age difference in basal glycolysis and glycolytic capacity (Supplemental Figure 12A-D). In young neutrophils, male neutrophils show higher basal glycolysis compared to female neutrophils with PMA treatment (p=0.028), pan-JAK inhibition (p=0.0163), and JAK2 inhibition (p=0.0025; Supplemental Figure 12E). No sex differences were observed in basal glycolysis of the aged neutrophils (Supplemental Figure 12E). In addition, young male neutrophils show an increase in glycolytic capacity compared to young female neutrophils under control conditions (p=0.0352), PMA treatment (p=0.0189), pan-JAK inhibition (p=0.0114), and JAK2 inhibition (p=0.004; Supplemental Figure 12F). There were no sex differences in glycolytic capacity of the aged neutrophils (Supplemental Figure 12F). Overall, our data show that JAK activity plays a direct modulatory role on neutrophil metabolism. This modulation differs across age, with young neutrophils dependent on JAK activation to increase glucose metabolism and PPP, while aged neutrophils use JAK2 activation to inhibit the breakdown of glycogen stores within the cell.

## Discussion

The results of this study elucidate specific intracellular pathways in neutrophils that are dependent on JAK activation. Prior to this study, the understanding of how JAK signaling regulates neutrophil function at the molecular level was limited. It is established that JAK2 plays a key role in neutrophil migration, primarily signaling through activated STAT3 in response to IL-8 stimulation [22]. However, the specific downstream proteins affected by JAK2 activation that directly drive cell motility remained largely unknown. Our findings reveal a novel and direct role of JAK2 regulation of neutrophil migration through the modulation of membrane composition and cytoskeletal remodeling (Figure 7A). Neutrophil migration is the directed movement of neutrophils guided by chemotactic signals such as chemokines or cytokines. This is not to be confused with infiltration, which refers to the process by which neutrophils exit the bloodstream and penetrate tissue, involving adhesion, transmigration through the endothelium, and subsequent tissue navigation. Our data shows that in both young and aged neutrophils, JAK2 activation inhibits the release of various inflammatory factors, IL-1α and VEGF, that influence neutrophil tissue infiltration [41, 72]. In both young and aged neutrophils, JAK2 activation modulates lipid incorporation into the plasma membrane and membrane stiffness by negatively regulating DHCR24 signaling, a key enzyme in cholesterol biosynthesis. JAK2 activation inhibits the release of SDF1α (CXCL12) by young neutrophils, a chemokine that typically facilitates their mobilization from the bone marrow into circulation. SDF1α is primarily expressed extracellularly by stromal cells within the bone marrow, where it plays a crucial role in neutrophil retention and release [73, 74]. Our data suggests that, in an autocrine fashion, neutrophils may regulate their retention in tissue by releasing SDF1α in the extracellular space, as previously shown in macrophages [75]. Additionally, SDF-1α has been shown to be a chemoattractant for neutrophils in vitro [22]. In aged neutrophils, JAK2 activation leads to an increase release of CXCL2, a key neutrophil chemoattractant. JAK2 does not enhance MARCKS activity in young or aged neutrophils, despite its role in facilitating ARP2/3 complex-mediated actin polymerization and branching, which is essential for pseudopodia formation and motility. However, JAK2 promotes the presence of ARPC2 and ARPC3, two critical ARP2/3 subunits, with ARPC2 serving as the nucleation point for actin polymerization and ARPC3 stabilizing the complex core, ensuring efficient cytoskeletal remodeling. Our data also shows that all JAK proteins may play a role in this cellular function, as shown by pan-JAK inhibition decrease of migration. Interestingly, most of the proteins modulated by pan-JAK inhibition are also modulated by JAK2 inhibition. Overall, these data demonstrate an age-dependent shift in migration regulation.

**Figure 7:**
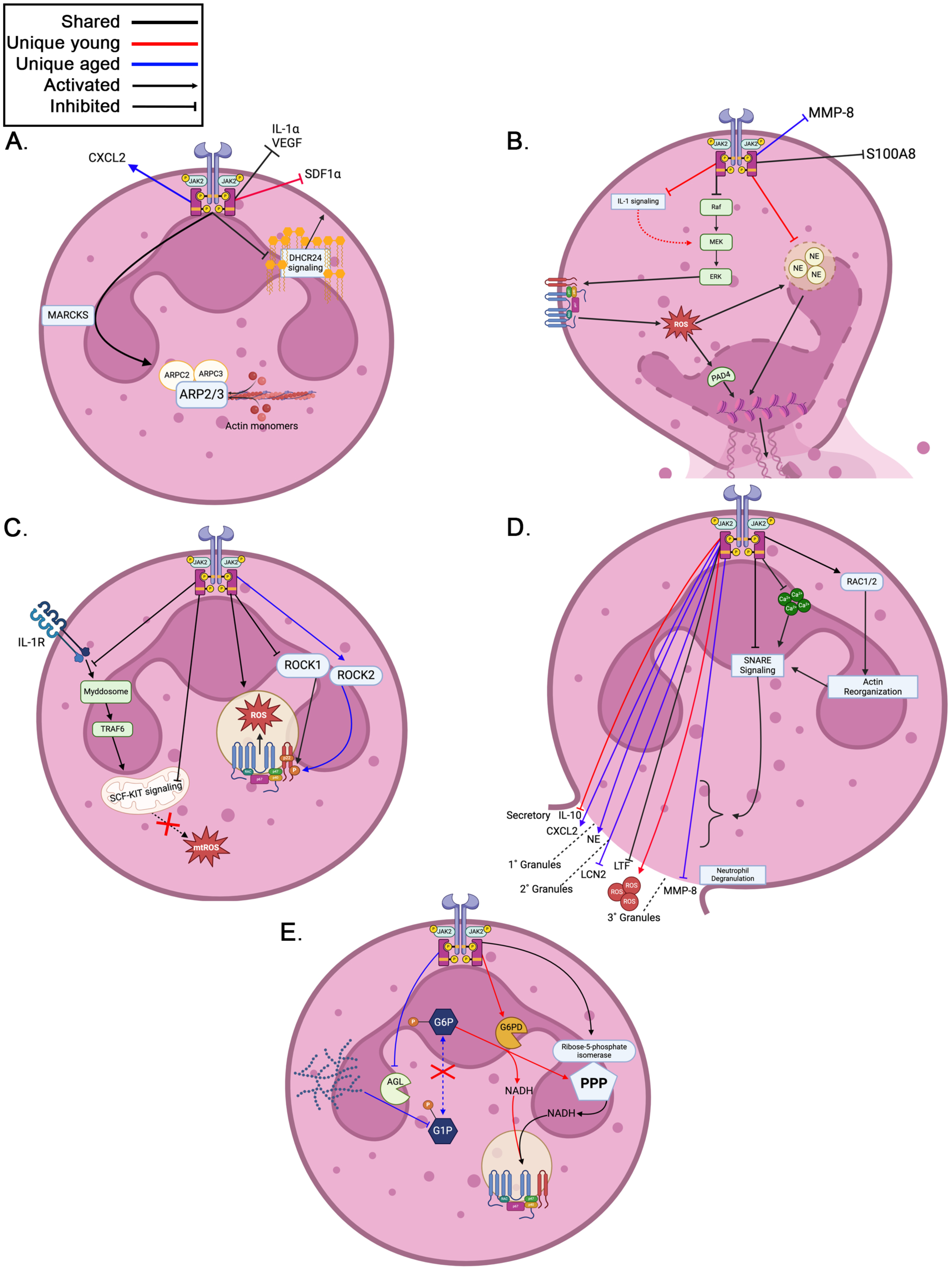
JAK2 differentially regulates neutrophil functions between young and aged neutrophils. **A.** JAK2 regulates neutrophil migration by modulating membrane composition via DHCR24 signaling, cytoskeletal remodeling through MARCKS and ARP2/3 complex, and differential secretion of chemokines such as CXCL2 and SDF1α. **B.** JAK2 indirectly influences NETosis in young neutrophils by modulating IL-1 signaling and NE activity, in both age groups JAK2 influences the secretion of NET-associated proteins MMP-8 and S100A8. **C.** JAK2 impacts ROS production through regulation of SCF-KIT and IL-1 signaling pathways and modulates key cytoskeletal regulators ROCK1/2, although in aged neutrophils JAK2 preferentially activates ROCK2. **D.** JAK2 regulates ROS-associated degranulation by modulating RAC1/2 and SNARE signaling pathways, affecting actin reorganization and the release of granule proteins such as MMP-8, LTF, LCN2, CXCL2, IL-10, and NE. **E.** JAK2 modulates glycolytic metabolism by promoting glucose shuttling into the pentose phosphate pathway through G6PD and ribose-5-phosphate isomerase in young neutrophils, while regulating glycogen breakdown through glycogen phosphorylase in aged neutrophils.

JAK regulation of NETosis is unclear, there is limited direct evidence of JAK regulation of key proteins that regulate this function. The treatment of neutrophils with tofacitinib, a JAK1/3 inhibitor with a lesser effect on JAK2, reduced NETosis in an ex vivo mouse model of lupus [76], no proposed mechanism was described in the study. NETosis occurs through chromatin decondensation and the release of decondensed DNA coated with granule proteins, a process regulated by MPO [77], NE [78], PAD4 [42], and cathepsin G [43]. Our findings indicate that JAK2 activation does not directly regulate NET formation. Further demonstrated by no change in PAD4, the master regulator of NETosis, across all conditions (Figure 3C). However, NETosis-related signaling pathways, including NE levels and IL-1 signaling, were affected by JAK inhibition in an age-dependent manner (Figure 7B). Our data also suggest that JAK2 activation can indirectly regulate NETosis through its modulation of ROS production via IL-1 signaling. ROS dependent NETosis is the most described form of NETosis. Previous studies have shown that IL-1*α* secretion coincides with ROS production; however, a direct link has not been established in neutrophils, though it has been observed in other cell types [79]. Additionally, IL-1β activates the PI3K-AKT pathway, which is used in neutrophil ROS production [80, 81]. ROS production is a critical driver of NET formation through its effect on MPO activity which leads to the activation of NE. NE, in conjunction with PAD4, promotes the citrullination of histones and nuclear envelope breakdown, essential steps in NET formation. Furthermore, JAK2 suppresses the release of markers of NETosis, such as, S100A8, a cytosolic protein, in both young and aged neutrophils, while selectively inhibiting the release of MMP-8, a tertiary granule-associated enzyme, in aged neutrophils exclusively. Overall, while JAK2 does not directly regulate NETosis, it influences this process indirectly by modulating ROS production through IL-1 signaling and suppressing the release of NET-associated markers, such as S100A8 and MMP-8, in an age-dependent manner.

There are conflicting data on JAK’s modulation of neutrophil ROS production. JAK inhibition (Baricitnib and Tofacitinib) showed no effect on the functional output of ROS production in human patients with rheumatoid arthritis [53]. However, JAK1-3 inhibitor (LAS194046) suppressed ROS generation in neutrophils from patients with asthma and COPD [54]. Our functional analysis showed no effect of JAK activity in neutrophil ROS production (Figure 4E-G). However, our data indicated that JAK2 may regulate key proteins in the assembly of the NADPH oxidase (NOX) complex formation, a key protein in neutrophil ROS production (Figure 7C). In young and aged neutrophils, JAK2 activation downregulates IL-1 signaling, shown to initiate the assembly of the Myddosome (MyD88, IRAK4, IRAK2) [82]. The Myddosome then will signal through TRAF6-ECSIT complex to activate mitochondrial ROS production [83]. JAK2 inhibits ROCK1 and upregulates the NOX complex in both young and aged neutrophils. Additionally, in young and aged neutrophils, JAK2 activation inhibits SCF-KIT signaling, ROS and NOS production, while simultaneously increasing the expression of NOX1. However, a notable difference, is that in aged neutrophils JAK2 activates ROCK2 where in young neutrophils ROCK2 is inhibited. ROCK1 and ROCK2 activate p22phox, a critical protein in the activation of the NOX complex. Our data suggest that JAK2 activation serves as a priming step in neutrophil ROS production by inhibiting mitochondrial ROS generation and promoting the assembly of the NOX complex. This facilitates the production of ROS upon further stimulation in both young and aged neutrophils.

JAK’s role in neutrophil degranulation is largely understudied. With one paper citing that IFN-*γ* (which signals through JAK1/2) signaling and neutrophil degranulation are responsible for the viral response in lungs of macaques infected with SARS-CoV-2, but there was no direct mechanism attributed to this [84]. Other studies have linked JAK1/2 activity to mast cell degranulation in response to inflammatory stimuli [85]. Even in the context of mast cells, the exact molecular mechanisms by which JAK modulates degranulation is unclear. Neutrophil degranulation is a tightly regulated process involving the exocytosis of granule content. This process is mediated by calcium influx, actin cytoskeletal reorganization, and SNARE-mediated membrane fusion [59]. Our findings demonstrate that JAK2 activation plays a distinct role in regulating neutrophil degranulation in both young and aged neutrophils (Figure 7D). Overall, JAK2 activation regulates degranulation by modulating calcium signaling, SNARE-mediated granule fusion, and actin reorganization. In young neutrophils, these modulations lead to the suppression of IL-10 and LTF release and the active release of ROS. On the other hand, aged neutrophils suppress MMP-8, LCN2, and LTF release, while simultaneously increasing the release of NE and CXCL2. These changes seem to be mediated by the activation of RAC1/2 signaling, which facilitates actin remodeling. In both young and aged neutrophils, JAK2 activation suppresses SNARE signaling, decreasing the fusion of secondary and tertiary granules to the plasma membrane. Interestingly, JAK2 induces the release of the chemokine CXCL2 (aged) and ROS (young) suggesting a positive regulation of the release of secretory and specific secondary vesicle content, respectively. Our data highlights an age-dependent shift in the mechanisms underlying neutrophil degranulation.

Neutrophils primarily rely on glycolysis for energy production. A recent study demonstrated that, in neutrophils, glucose can be diverted from the glycolysis into the PPP to generate NADPH and facilitate ROS production, a process dubbed glucose shuffling [68, 69]. Overall, our findings reveal that JAK2 directly modulates neutrophil metabolism by regulating glycogen metabolism in an age dependent manner (Figure 7E). In young neutrophils, JAK2 activation leads to the activation of glucose-6-phosphate dehydrogenase (G6PD) which, with ribose-5-phosphate isomerase move glucose into the PPP cycle to generate NAD+. In aged neutrophils, JAK2 activation modulates the breakdown of glycogen into glucose-1-phosphate thereby increasing the availability of glucose. Amongst immune cells, neutrophils contain the most glycogen stores, whether these stores are influenced by age still needs to be determined [86]. This may indicate that young neutrophils may preferentially prioritize ROS generation over pure energy demands. These data suggest that JAK2 regulation of glucose metabolism changes in neutrophils with age, JAK2 activation in young neutrophils preferentially lead to pathways to increase ROS production, JAK2 activation in aged neutrophils leads to decrease breakdown of glycogen. It remains unclear from our data whether the metabolic shift in aged neutrophils is driven by a decline in glucose uptake.

## Conclusion

This study provides new mechanistic insights into JAK modulation of neutrophil function. This modulation is influenced by age and not sex. Our data show that, in young and aged neutrophils, JAK2 activation directly modulates migration through membrane composition or cytoskeletal rearrangement. On the other hand, JAK2 activation inhibits release of chemokine in young neutrophils, while increase its release in aged neutrophils. Our findings indicate that JAK2 activation does not directly regulate NET formation, though it may play an indirect role by modulating ROS production. Additionally, our data demonstrate that JAK2 activation differentially regulates neutrophil degranulation, with young neutrophils exhibiting RAC1/2-mediated actin remodeling and aged neutrophils showing impaired granule release. Finally, JAK2 signaling promotes PPP activity in young neutrophils while decreasing glycogen breakdown in aged neutrophils.

This study has several limitations. This includes the fact that neutrophils were extracted from the bone marrow, as such we are probing this cellular mechanism in naïve cells. We acknowledge that it is possible that exiting the bone marrow may change the effect of JAK2 signaling on these cells. Additionally, the extracted cells likely represent a mix of immature, mature (marginated), and aged neutrophils. We postulate that most of the cells in our isolation are mature marginated neutrophils. Another limitation to note is that this study is focused on mouse neutrophils. Whether these molecular mechanisms are also present in human neutrophils will have to be confirmed in subsequent studies.

However, with mice being the preferred animal to model human disease, we believe it is important to fully understand how these cells are regulated to better interpret future findings. Lastly, the PMA concentration used in this study was lower than the concentration typically used in other published work. This approach was chosen to achieve effective activation while minimizing excessive cell death. To conclude, despite these limitations, this study provides a foundational understanding of how JAK2 modulates downstream effector proteins and regulates neutrophil function across biological aging, offering critical insights that may inform future therapeutic strategies targeting immune dysfunction in aging and inflammatory diseases.

## Materials and methods

### Animals

Young (3 months of age) and aged (22+ months of age) wildtype (C57Bl6/J) mice were generated in our animal facility or purchased from Jackson Laboratory. Functional assays were conducted using 4–6 mice per sex/age group. Each mouse provided a sufficient number of cells to be allocated across all experimental conditions, enabling their use in functional assays and mass spectrometry analysis. Mice were housed under normal conditions, a 12:12 hour light dark, and had access to food and water ad libidum. All animal procedures were approved by the Institutional Animal Care and Use Commitee at West Virginia University.

### Neutrophil isolation

Mice were euthanized by isoflurane overdose, and bone marrow collected using sterile RPMI solution with 2mM EDTA. Neutrophils were isolated from the cell suspension using the negative selection Biolegend MojoSort Kit (Cat# 480058). Protease and phosphatase inhibitor cocktail (ThermoFisher Cat# 78442) was added during the isolation process to minimize protein degradation. This isolation yields on average 3.8x10^6^ cells/mL in 4 mL of media. Flow analysis shows an 80% neutrophil enrichment post-isolation, compared to the pre-isolation of 40% neutrophils (Supplemental Figure 1A).

### Culture preparation and collection for functional and proteomic analysis

For ROS and NET functional assays, isolated neutrophils were plated on a poly-D-Lysine coated glass bottom petri dish with RPMI media at a density of 2.5x10^5^ cells/mL. For the migration assay, glycolysis assay, and proteomic collection plates were not treated with Poly-D-Lysine. Cultures were placed in the cell culture incubator (at 37 degrees Celsius with 95% humidity and 5% CO_2_) for 30 min. Inhibitors (1µM; Baricitinib-Selleckchem Cat# S2851 or AZD1480-AdooQ Cat# A10110) were added to cultures for 1 hour, followed by the addition of Phorbol 12-myristate 13-acetate (PMA-20ng/mL; Sigma-Aldrich Cat# 524400) for an additional 45 min, prior to collection (except for the migration assay). For proteomic collection, cultures were transferred to FACS tubes, spun at 300g for 10 minutes, supernatants and cell pellets were collected, and a protease and phosphatase inhibitor cocktail was added. These samples were flash frozen and stored until sent to IDeA Proteomics at the University of Arkansas Medical Sciences for mass spectrometry.

### Functional assays

#### Migration assay

Boyden chamber insets, with 3µm pores, were purchased from Abcam (Cat# ab235692). These transwell chambers were placed on a 96 well plate, the lower chamber of the well contained 150µL of media and the chemoattractant Interleukin 8 (20ng/mL; Kingfisher BioTech Cat# RP0357H-005). 5x10^4^ neutrophils were added to the top chamber of each transwell in 100µL and PMA (20ng/mL) was added to each well. Cultures were then left to incubate for 2 hours in the incubator. At the end of the incubation period, the number of cells in the lower chamber was counted using the Countess 3 automated cell counter (FisherScientific).

#### Modified CellROX assay

Isolated neutrophils were plated on a poly-d-lysine coated glass bottom petri dish. 2.5x10^5^ neutrophils were added for the Reactive Oxygen Species (ROS) production/degranulation assays. CellROX green reagent (5µM; ThermoFisher Cat# C10444) was applied to the cells at the same time as the PMA application (after the inhibitor incubation as stated above). After 45 minutes, plates were collected, aspirated, and fixed with 4% paraformaldehyde (Electron Microscopy Sciences Cat#30525-89-4) for 1 hour. Cells were counterstained with DAPI (ThermoScientific Cat# 62248) and Phalloidin (ThermoFisher Cat# A22287). Plates were blinded for image analysis and quantification. 3 images were obtained per plate using the epifluorescent Echo Revolve (Echo). For quantification purposes, cells that had higher green fluorescence within the phalloidin cell boundaries were counted as cells upregulating ROS production (Figure 4A, red arrow); and those with green fluorescence immediately adjacent to the phalloidin cell boundary were counted as degranulated cells (Figure 4A, yellow arrow). All ROS data was normalized by the total number of DAPI nuclei in each image prior to analysis.

#### Neutrophil extracellular trap (NET) analysis

Isolated neutrophils were plated on poly-d-lysine coated glass bottom petri dishes. 2.5x10^5^ isolated neutrophils were added and incubated with the inhibitor and then PMA as stated above. After incubation, cells were fixed with 4% PFA, and stained with DAPI, anti-Histone 1 (ThermoFisher Cat#3005-MSM7-P1), and anti-Myeloperoxidase (Proteintech Cat# 22225-1-AP). Plates were blinded for image analysis and quantification. 3 images were obtained per plate using the epifluorescent Echo Revolve. NETs were characterized as Histone 1 positive, DAPI positive, and MPO positive (Figure 3A, H1 positive cells). Total amount of NET positive data was divided by total DAPI nuclei for normalization, then analyzed.

### Mass spectrometry

Frozen supernatants (secretome) and cell pellet (cell fraction) samples were sent to the IDeA National Resource for Quantitative Proteomics for processing and analysis. The acquired proteomics data were post-processed and grouped based on condition, sex, and age (n= 4 per group). Data normalization was performed using a Loess smoothing model, which employs locally weighted regression to estimate and correct for bias. Contrast comparison values, including log fold changes, FDR-adjusted p-values, and average intensities, were obtained from IDeA. Subsequently, in-house analyses were conducted using Ingenuity Pathway Analysis (IPA) to examine differences across conditions, age groups, and sexes. Proteins that fell below the detection threshold in some samples were excluded from the entire group to maintain a consistent sample size of n=4, ensuring statistical comparison. All comparisons were normalized to age- and sex-matched controls.

### Functional glycolysis (metabolic) assay

Neutrophils were isolated and incubated with the inhibitors as described above. Cells were resuspended in Seahorse XF RPMI media, supplemented with 2 mM glutamine, and seeded at a density of 2x10^5^ cells/well at 50 uL in XF96-well plates (Agilent Cat#103794-100) coated in Cell-Tak adhesive (Corning Cat# 354240). Plates were then centrifuged at 200xg for 1 minute and allowed to incubate in a non-CO2 incubator. Plates were then analyzed by the Seahorse Bioscience XF Pro instrument, which measures the extracellular acidification rate over time. Sequential injections of glucose, oligomycin, and 2-Deoxy-D-glucose (2-DG) were injected at 10 mM, 1.5 uM, 100 mM respectively. Assay runs for a length of around an hour and forty-five mins.

### Luminex (ELISA)

22-plex Luminex assays (Bio-techne, CUSTOM-LXSA-M22) were run using supernatants collected for mass spectrometry. The proteins measured were pro-inflammatory cytokines (IL-1β, IL-1α, IL-6, IL-17, TNF-α, IFN-*γ*), anti-inflammatory cytokines (IL-10), chemokines (CCL2, CXCL1, CXCL2, CXCL10, SDF1α), growth factors (G-CSF, IGF-1, VEGF), proteases (S100A8, Serpin E1, MMP-8, MMP-9) and complement receptor (C1qR). Luminex assays were run according to manufacturer’s protocol by the Flow Cytometry and Single Cell core facility at West Virginia University Health Science Center. Assays were run on the Luminex MAG PiX instrument using the xPONENT software version 4.3 update 1.

### Statistical Analysis

Analyses were normalized to the control (which has no PMA and no inhibitors) to generate a value indicative of change from the control condition for all data (except if raw data is indicated). Data from blinded unbiased stereological counts and migration assays were statistically analyzed using One-Way or Two-Way Analysis of Variance (ANOVA), evaluating interactions between condition, age, and/or sex. For the metabolism assay, raw data were collected, and the Area Under the Curve (AUC) was calculated for each group. AUC values were normalized to their respective controls and compared using One-Way or Two-Way ANOVA, followed by Tukey’s post-hoc analysis. Mass spectrometry data were analyzed by generating functional pathway and protein-level heat maps, with positive and negative z-scores indicating up- and down-regulation, respectively. Fisher’s exact test was employed as a post-hoc analysis for heatmaps. For individual protein analyses, intensity values from each animal were subjected to Two-Way ANOVA to assess the effects of condition and age on protein abundance, with Tukey’s post-hoc tests used to identify significant group differences. All data are presented as mean ± standard error of the mean (SEM), with statistical significance set at p < 0.05. Statistical analyses were conducted using GraphPad Prism 10.4 (Dotmatics, San Diego, CA).

## Acknowledgements

We want to thank everyone in Dr. Aminata Coulibaly’s lab who aided with the work seen throughout this paper. We gratefully acknowledge the OLAR staff at West Virginia University for their dedicated care of our animals. We extend our sincere thanks to Dr. Hollander and Ethan Meadows at WVU Mitochondria functional assessment core for their invaluable assistance with the Seahorse assays. We also thank the IDeA Proteomics Core at the University of Arkansas for Medical Sciences for processing our mass spectrometry samples and providing expert analysis support. Finally, we are grateful to Dr. Kathy Brundage and the WVU Flow Cytometry Core for their guidance and assistance with flow cytometry experiments.

## Author contributions

JWF worked with APC to conceptualize the study, wrote and edited the manuscript. JWF conducted, collected, and analyzed all experiments. MK performed all Seahorse assays and contributed to writing of the manuscript. FM, EH, and NS quantified microscopy images for NETosis, ROS production, and degranulation analyses. CW acquired and managed all animals used in the study. APC provided the framework and concept for the study, provided supervision and oversight throughout the project, and edited and finalized the manuscript.

---

**Supplemental Figure 1:**
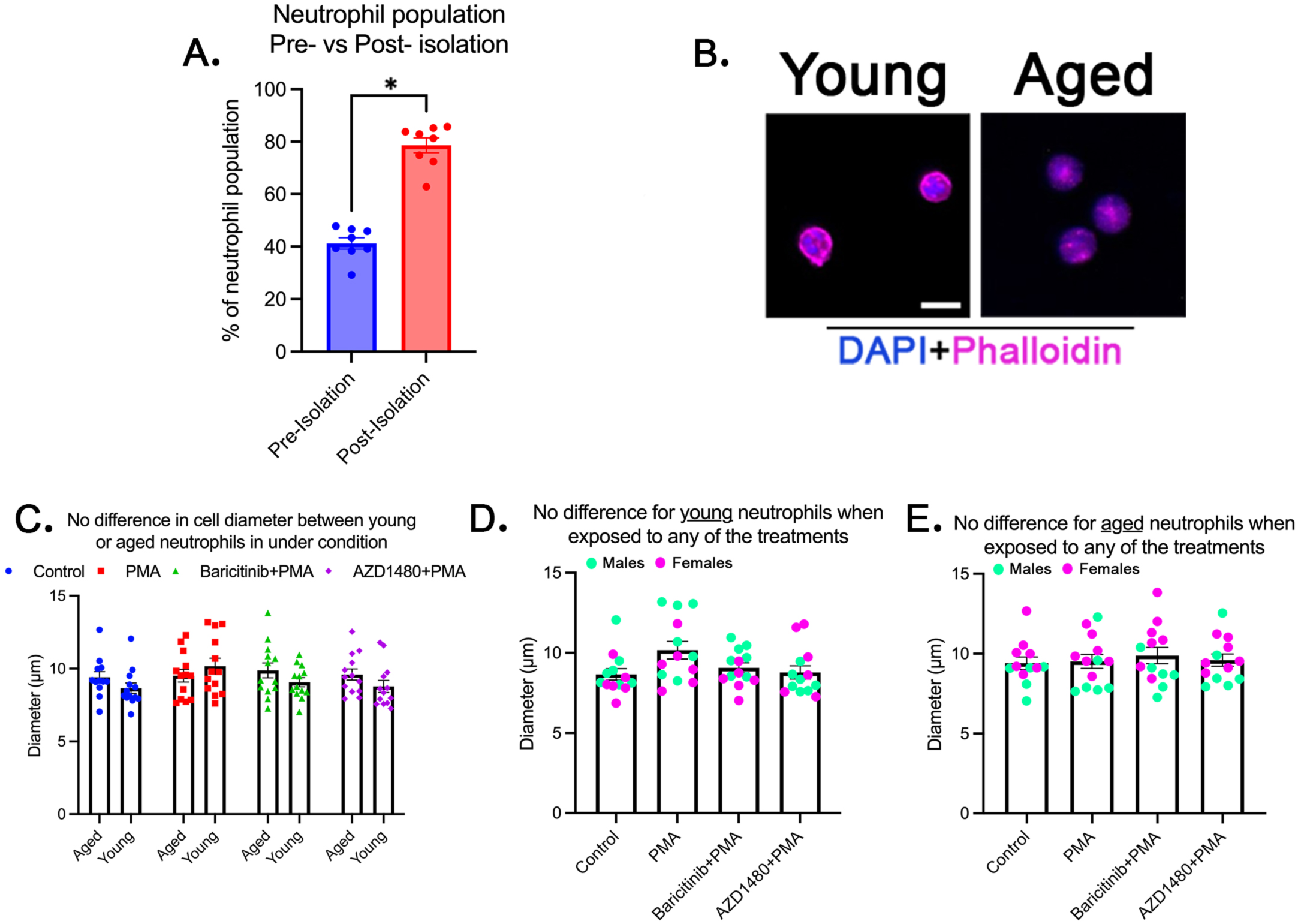
Neutrophil isolation successfully enriches neutrophils in the cell suspension, with no observed differences in cell diameter across conditions, age groups, or sexes. **A.** Quantification of neutrophil population pre- and post-isolation, demonstrating a significant increase post-isolation (n=8). **B.** Representative fluorescent microscopy images of young and aged neutrophils stained with DAPI (nuclear stain) and phalloidin (actin cytoskeleton). **C.** Quantification of neutrophil diameter showing no significant difference between young and aged neutrophils under any condition (n=13/condition). **D-E.** No significant changes in neutrophil diameter in response to JAK inhibition across age groups (n=13/condition). p-value: * <0.05; bars represent mean ± SEM;

**Supplemental Figure 2:**
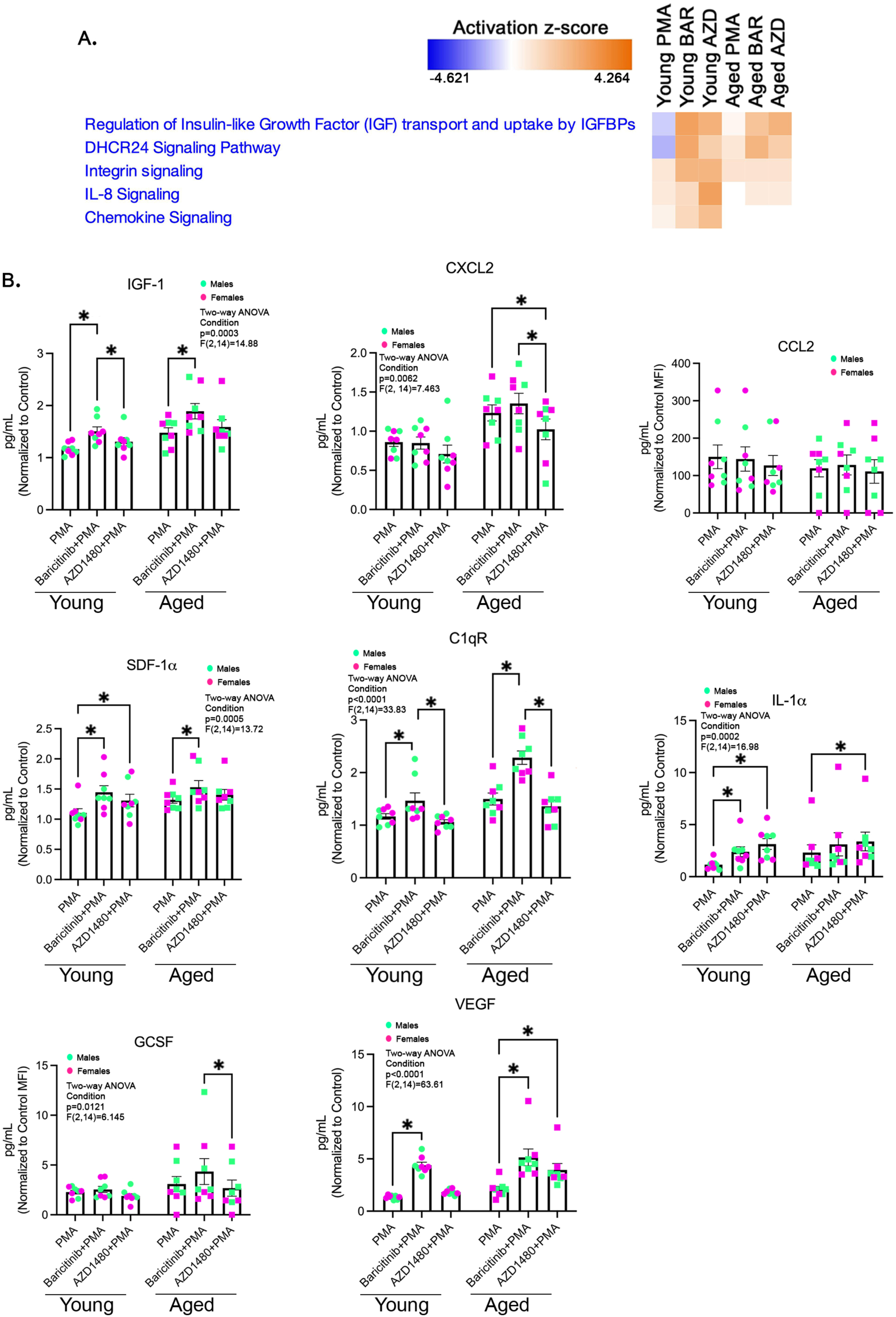
JAK modulates secretory proteins linked to neutrophil migration in an age-dependent manner. **A.** Functional heatmap of pathway activation z-scores highlighting changes in migration-associated signaling, in young and aged neutrophils across conditions (normalized to control). **B.** Luminex analysis quantifying the secretion of key migration-related cytokines and chemokines (IGF-1, CXCL2, CCL2, SDF-1α, C1qR, IL-1α, G-CSF, and VEGF) in young and aged neutrophils across different treatment conditions (normalized to control). Repeated Measures Two-way ANOVA, p-value: * <0.05; bars represent mean ± SEM; n=8/age/condition.

**Supplemental Figure 3:**
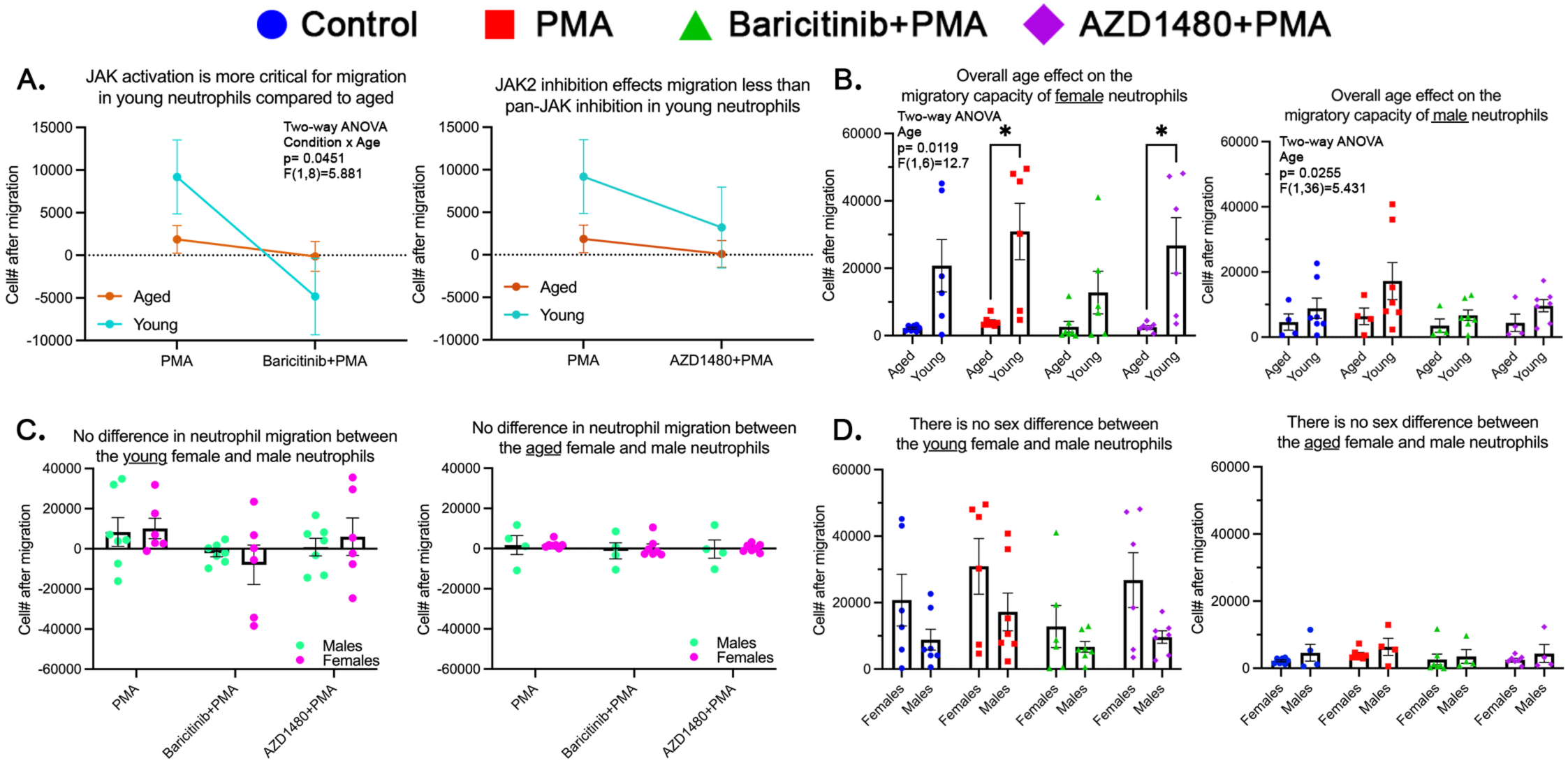
JAK regulates neutrophil migration in an age dependent manner, but is not sex-dependent. **A.** Migration differences between young and aged (combined sex and normalized to control) neutrophils under JAK inhibition (young: n=13/condition; aged: n=11/condition). **B.** Comparison of age-related effects on neutrophil migration, with either male (young: n=6/condition; aged: n=7/condition) or female (young: n=7/condition; aged: n=4/condition) neutrophils. **C.** No significant sex-based differences in neutrophil migration (normalized to control) were observed in young (female: n=7; male: n=6) or aged (female: n=4; male: n=7) neutrophils. **D.** Raw comparison revealed no significant sex-based differences in neutrophil migration in young or aged neutrophils. p-value: * <0.05; bars represent mean ± SEM;

**Supplemental Figure 4:**
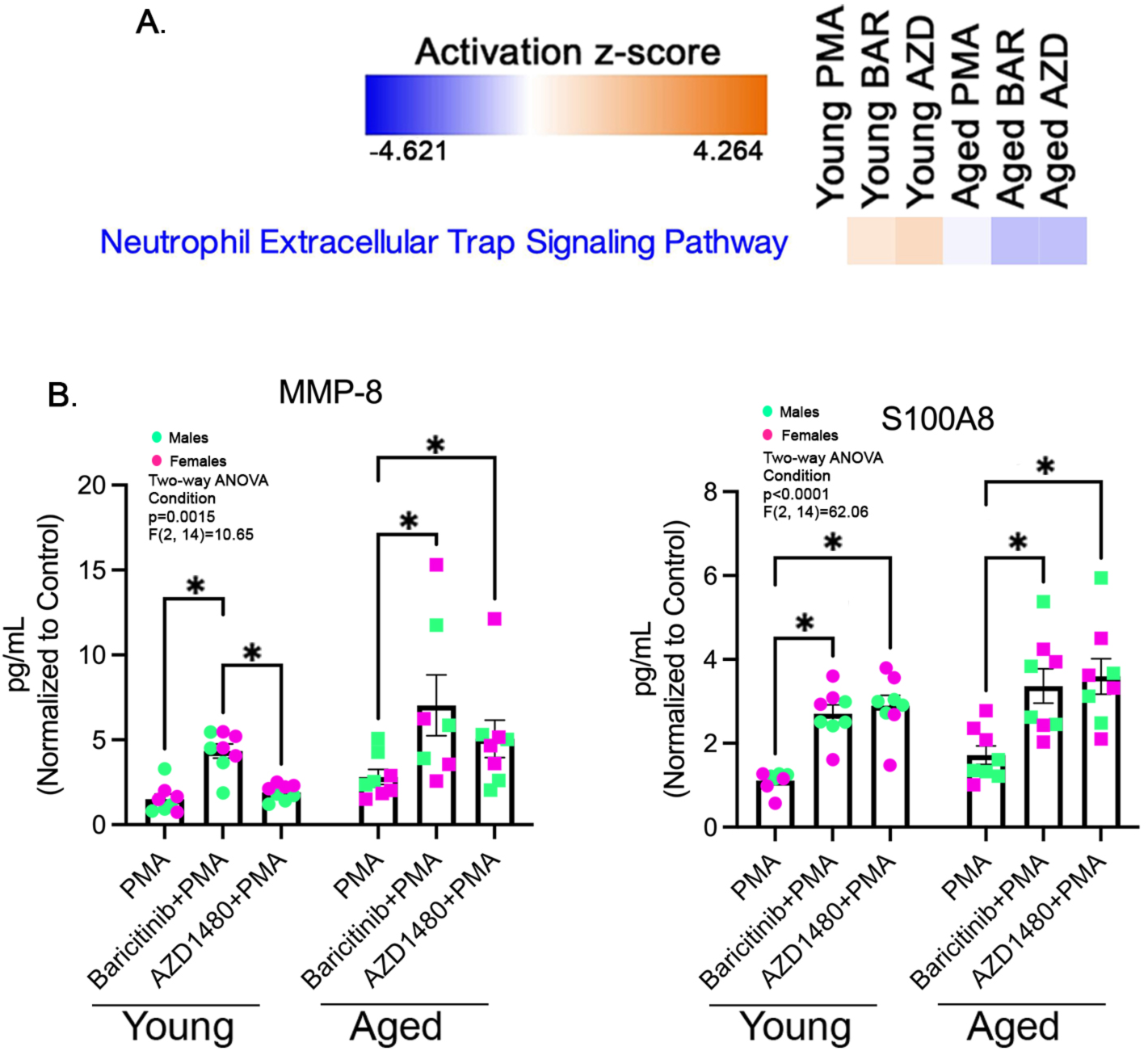
JAK inhibition does not affect NETosis-associated secretome but alters the composition of some of the secreted proteins. **A.** Activation z-score of functional heatmap showing NETosis pathway regulation (normalized to control). **B.** Luminex analysis of MMP-8 and S100A8 secretion in young and aged neutrophils following JAK inhibition (normalized to control; n=8/age/condition). p-value: * <0.05; bars represent mean ± SEM

**Supplemental Figure 5:**
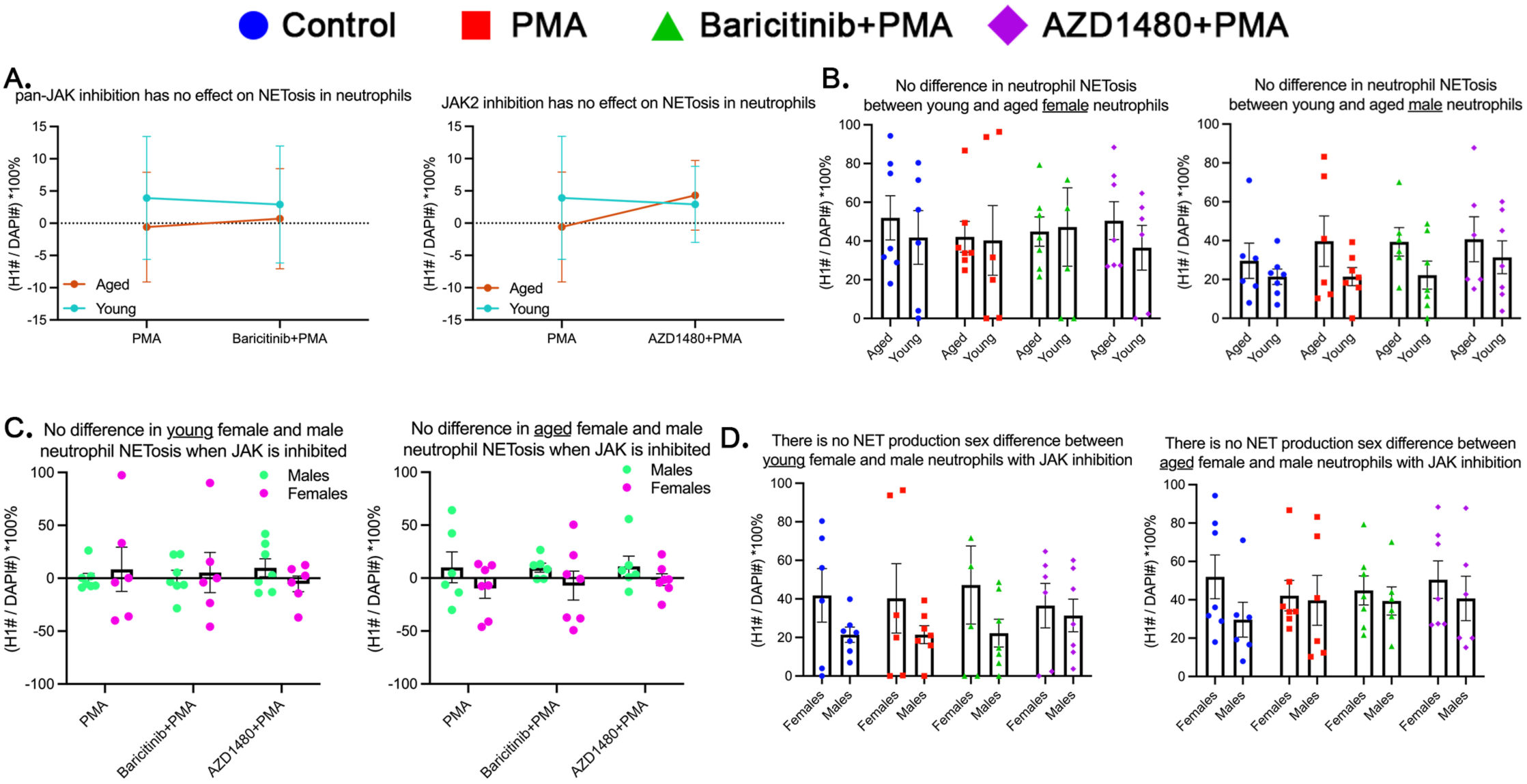
There is no age or sex differences in NETosis following JAK inhibition. **A.** No significant effect of JAK inhibition on NET formation in young (n=13/condition) or aged (n=13/condition) neutrophils (Repeated Measures Two-way ANOVA, p-value: * <0.05; bars represent mean ± SEM). **B.** No age-related differences in NETosis in young (female= 6; male=7) or aged (female= 7; male=6) neutrophils (Repeated Measures Two-way ANOVA, p-value: * <0.05; bars represent mean ± SEM). **C.** No significant sex-based differences in NETosis in young or aged neutrophils (Repeated Measures One-way ANOVA, p-value: * <0.05; bars represent mean ± SEM) normalized to control or **D.** in the raw comparison (Repeated Measures One-way ANOVA, p-value: * <0.05; bars represent mean ± SEM).

**Supplemental Figure 6:**
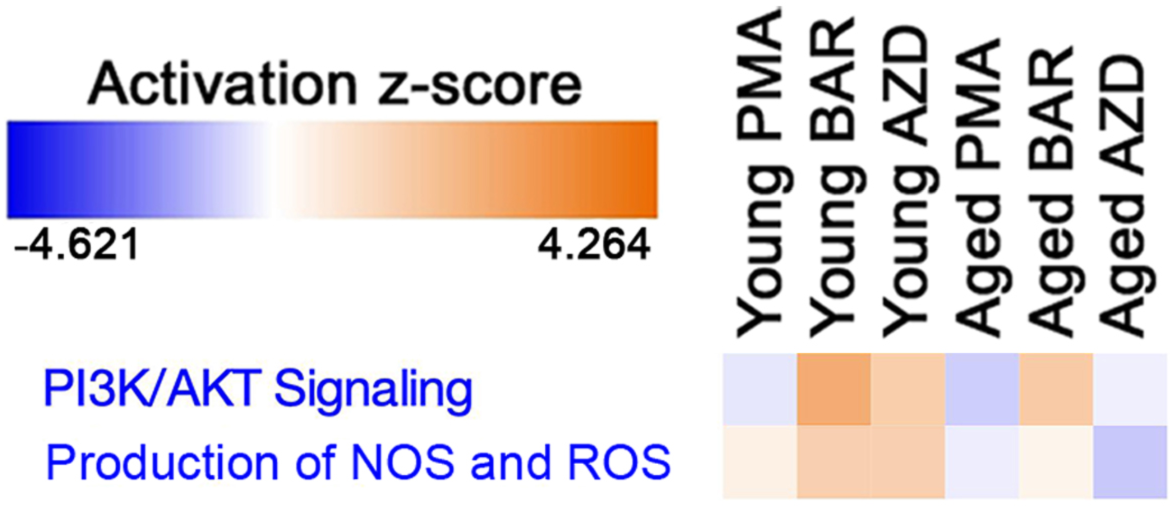
JAK signaling impact on ROS-related secretome pathways. Functional heatmap displays changes in PI3K/AKT and NOS/ROS production pathways following JAK inhibition across young and aged neutrophils (normalized to control).

**Supplemental Figure 7:**
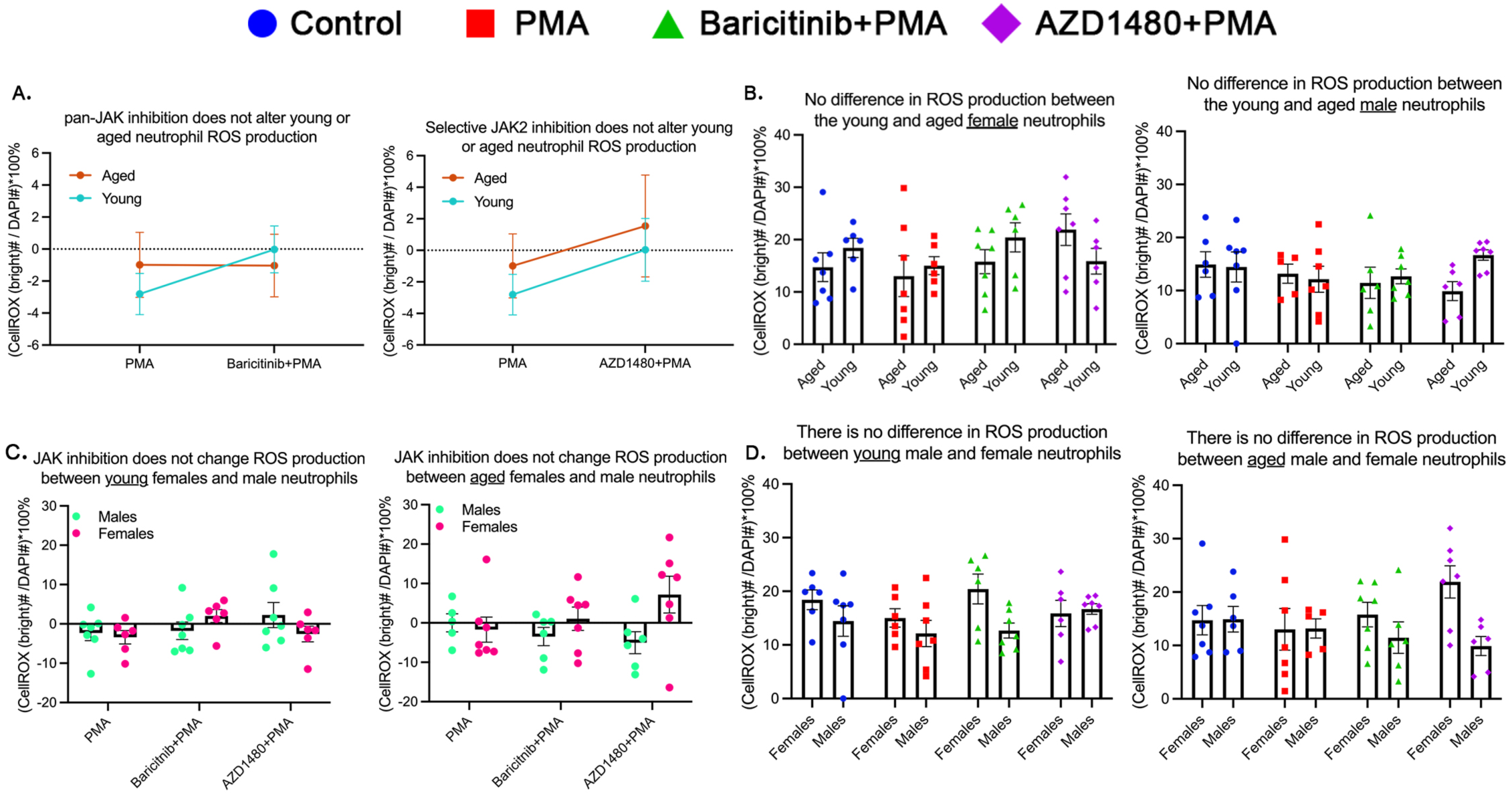
No age or sex differences in ROS production following JAK inhibition. **A.** No significant changes in ROS production with JAK inhibition in young (n=13/condition) or aged (n=13/condition) neutrophils. **B.** No age-related differences in raw ROS production comparison in young (female= 6; male=7) or aged (female= 7; male=6) neutrophils. **C.** No significant sex-based differences in ROS production in young or aged neutrophils normalized to control or **D.** in the raw comparison. p-value: * <0.05; bars represent mean ± SEM.

**Supplemental Figure 8:**
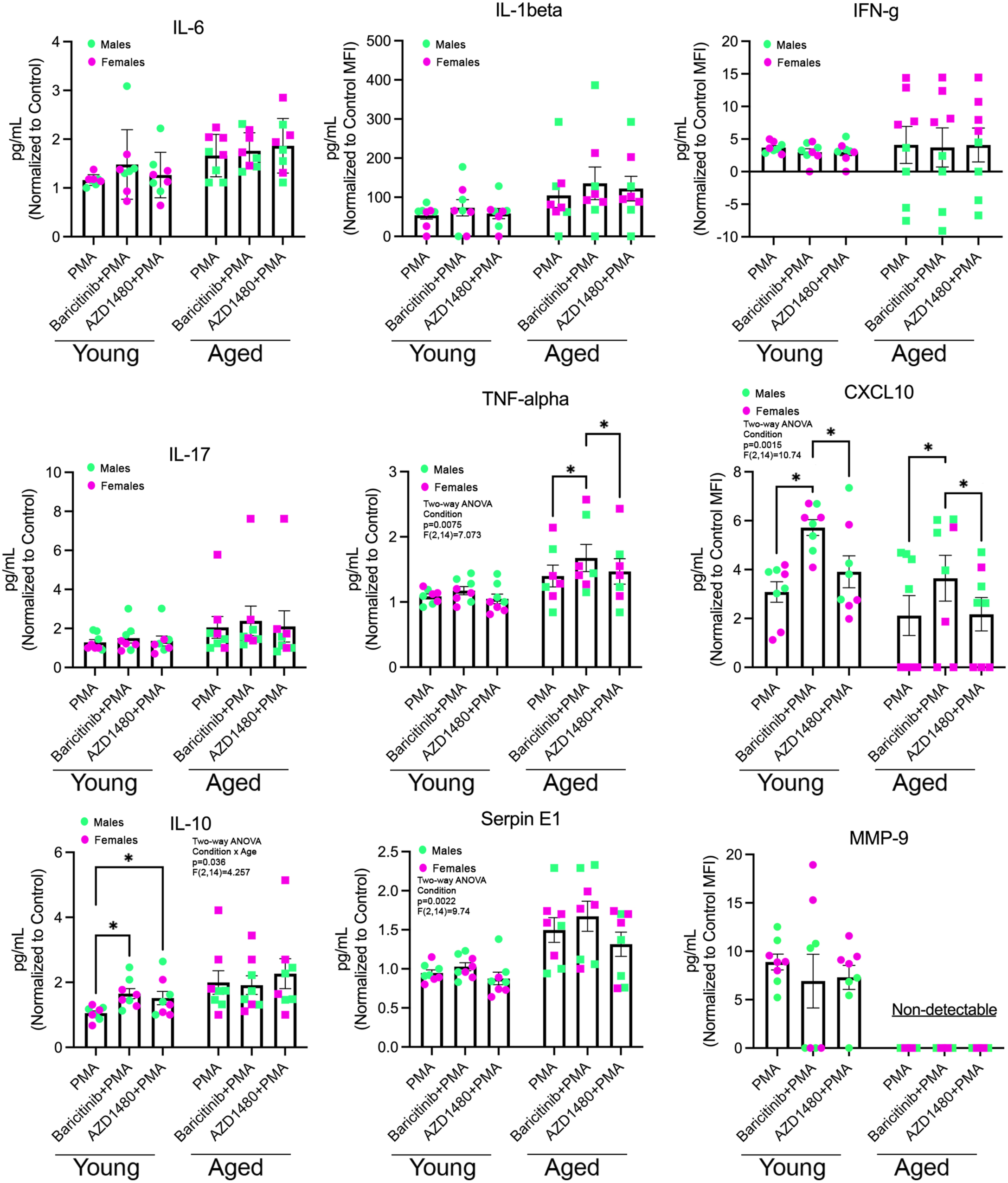
Luminex analysis of secreted proteins following JAK inhibition. IL-6, IL-1β, IFN-γ, IL-17, TNF-α, CXCL10, IL-10, Serpin E1, and MMP-9 expression comparisons. Normalized to control, p-value: * <0.05; bars represent mean ± SEM; n=8/age/condition

**Supplemental Figure 9:**
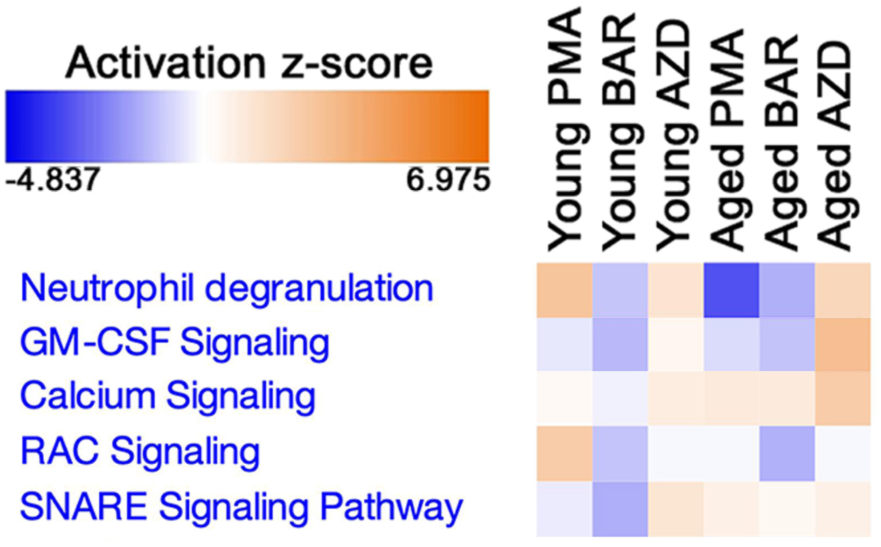
JAK-mediated regulation of neutrophil degranulation intracellular pathways. Heatmap illustrates changes in GM-CSF signaling, calcium signaling, RAC signaling, and SNARE-mediated membrane fusion across age and treatment conditions.

**Supplemental Figure 10:**
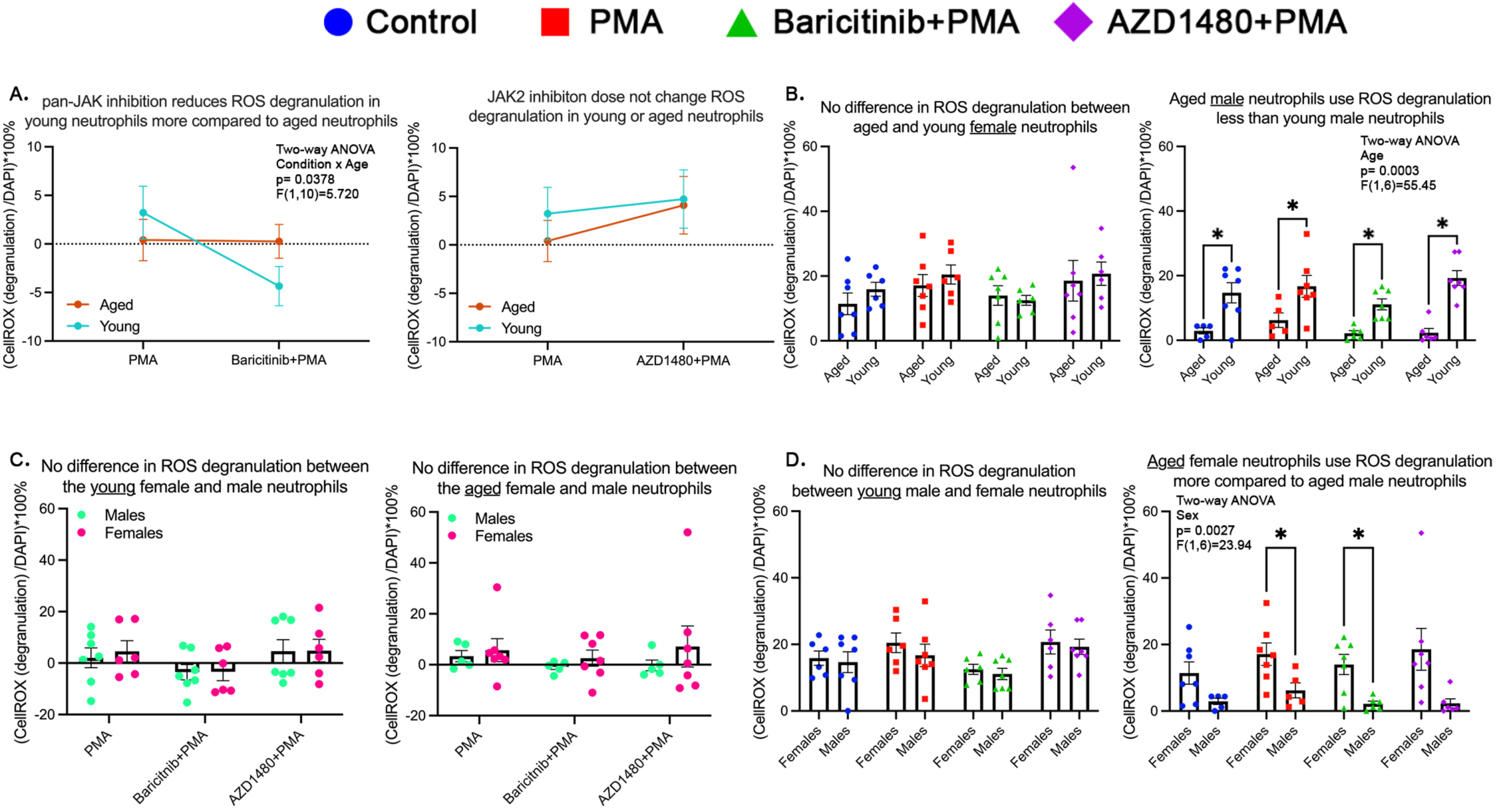
There are no observed sex differences in neutrophil degranulation across age groups; however, age-related differences in degranulation may be driven by females. **A.** Pan-JAK inhibition reduces ROS-associated degranulation in young (n=13/condition) but not aged (n=13/condition) neutrophils. **B.** Raw comparison, stratified by sex, shows young male (n=7/condition) neutrophils degranulate more than aged male (n=6/condition) neutrophils, but we do not see this trend in young (n=6/condition) and aged (n=7/condition) female neutrophils. **C.** No significant sex-based differences in degranulation in young or aged neutrophils, normalized to control**D.** However, in the raw comparison age neutrophils showed a sex difference, where male neutrophils degranulate less than female neutrophils, showing aged male neutrophils may be driving this age difference. p-value: * <0.05; bars represent mean ± SEM.

**Supplemental Figure 11:**
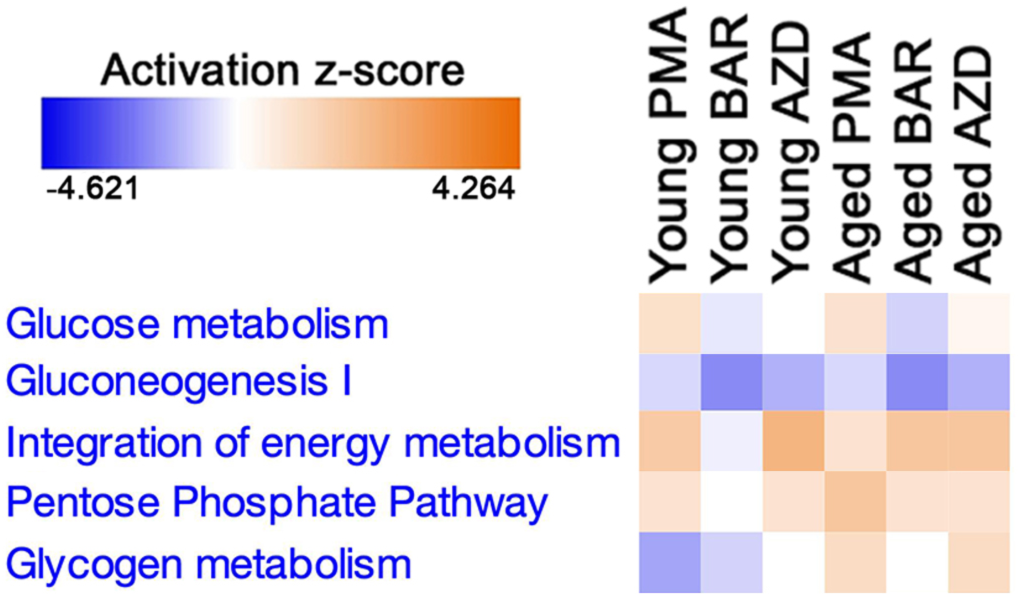
JAK signaling regulation of secretome metabolic pathways in neutrophils. Functional heatmap displays changes in glucose metabolism, gluconeogenesis, pentose phosphate pathway, and glycogen metabolism secretome across young and aged neutrophils following JAK inhibition.

**Supplemental Figure 12:**
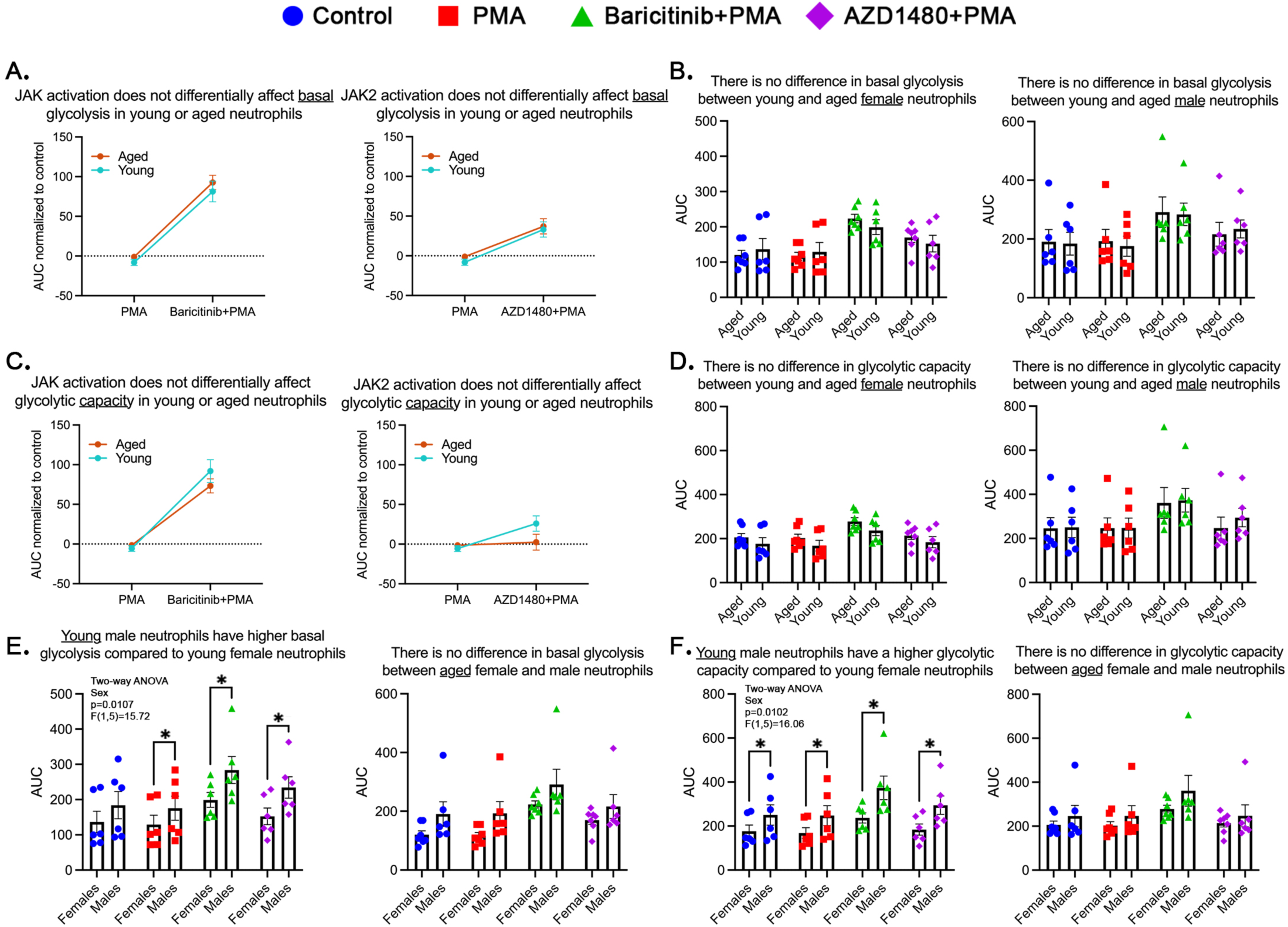
There are no age-dependent differences in neutrophil metabolism, however young neutrophils display a sex difference. **A.** JAK activation does not differentially affect basal glycolysis between ages, normalized to control (n=12/condition; aged: n=13/condition). **B.** Raw comparison shows no significant differences in basal glycolysis between young (female=6, male=6) and aged (female=6, male=7) neutrophils. **C.** Glycolytic capacity remains unchanged across age groups, normalized to control. **D.** Raw comparison shows no change in glycolytic capacity between ages. **E.** Young male neutrophils have a significantly higher basal glycolysis compared to young female neutrophils, but no sex differences in aged neutrophils. **F.** Young male neutrophils exhibit higher glycolytic capacity compared to young female neutrophils, but no sex differences in aged neutrophils. p-value: * <0.05; bars represent mean ± SEM.

